# Resolving ScRNA-Seq Signatures of Antigen-Specific CD4^+^ T Cells in Tolerance across Semi-Allogeneic Transplantation and Pregnancy

**DOI:** 10.1101/2025.07.06.663404

**Authors:** Michael S. Andrade, Grace E. Hynes, Zara R. Suran, Dengping Yin, Maria-Luisa Alegre, Peter T. Sage, Anita S. Chong

## Abstract

Transplantation of allogeneic organs requires lifelong immunosuppression to prevent rejection. Prior sensitization and resultant memory T cells are barriers to achieving successful transplant tolerance. In reproductive immunology by contrast, pregnancy represents a spontaneous model of tolerance where the semi-allogeneic fetus evades rejection even in multiparous or rejection-sensitized mothers. CD8^+^ T cell phenotypes of tolerance and rejection have been previously reported in transplant and pregnancy, but the transcriptional states of donor and fetus-specific CD4^+^ T cells remain poorly defined. Here, we performed Single-cell RNA-sequencing on endogenous, antigen-specific CD4^+^ T cells across models of allogeneic heart transplants and naïve or paternal skin-sensitized pregnancy. We identified expanded T follicular helper (Tfh) and non-follicular effectors in transplant rejection absent in tolerance. Naïve pregnancy resulted in a modest expansion of effector clusters with transcriptional quiescence that mirrored virgin mice. Successful sensitized pregnancy resulted in expanded Tfh clusters consistent with increased fetal-specific antibodies and limited non-Tfh effector responses. Most striking were the extensive changes imposed on donor-specific *Foxp3*^pos^ regulatory T cells (Tregs) resulting in the co-clustering together with *Foxp3*^neg^ T conventional cells (Tconvs) in transplant tolerance and the emergence of a *Foxp3*^neg^ Type I Regulatory cluster observed in pregnancy of sensitized dams. Finally, we showed that these transcriptomes were relevant and enriched in human datasets of health and disease respectively. Thus, the context-dependent signatures of antigen-specific CD4^+^ T cells provide new insights into their divergent responses to allogeneic conflict at the intersection of transplant and reproductive immunology.

**Graphical Abstract:** 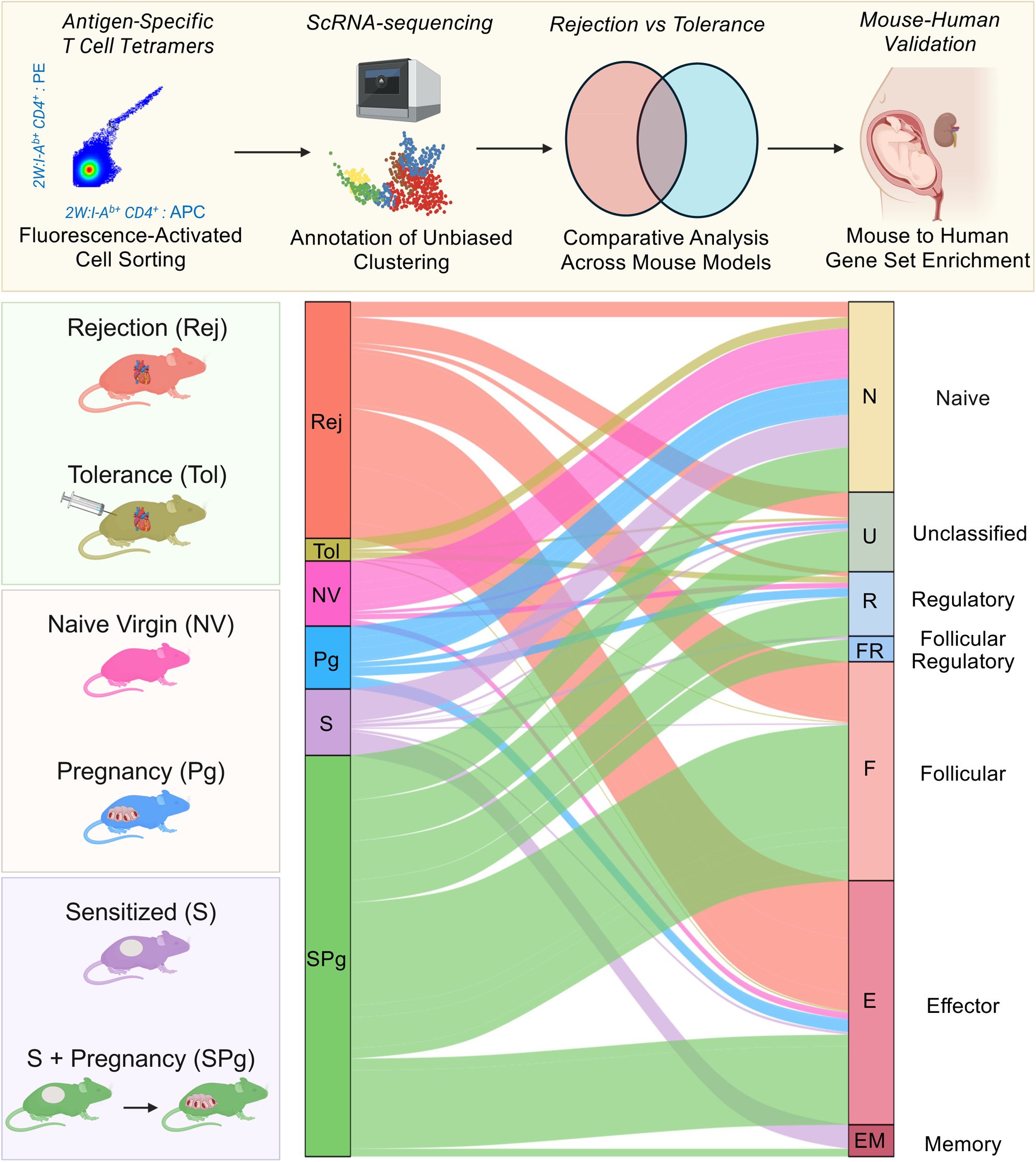

## INTRODUCTION

Immune tolerance is a state of hyporesponsiveness towards self or non-self antigens^1^. Transplanted allogeneic organs invariably trigger immune-mediated rejection, which can broadly be classified as T cell or antibody mediated, with both occurring in settings of inadequate immunosuppression ^2,3^. While pharmacological immunosuppression is the clinical standard of care to prevent rejection, the ability to induce lifelong transplantation tolerance through transient immunotherapy remains a central but elusive goal for the field. Multiple approaches, including co-stimulation blockade, successfully induce transplantation tolerance in rodent models, however, translating them into clinical practice has proved to be a significant challenge ^4^. High frequencies of alloreactive memory T and B cells represent a key barrier to the induction and maintenance of durable transplantation tolerance, and there is an absent conceptual framework for reprograming these populations towards transplantation tolerance ^5^.

In contrast to transplantation, pregnancy is recognized as a spontaneous model of immunological tolerance whereby the maternal immune system accommodates the semi-allogeneic fetus despite expression of paternal antigens. The divergent immune responses in immunologically competent adults to the semi-allogeneic fetus versus transplanted organs was noted as an immunological paradox by Medawar in 1953^6^. Multiple immune mechanisms contributing to tolerance of the semi-allogeneic fetus have been identified and classified as those acting locally at the maternal-fetal interface or systemically through the induction and expansion of fetus-specific regulatory T cell (Treg), CD8^+^ T cell exhaustion, and CD4^+^ T cell tolerance through interactions with immunomodulatory B cells ^7, 8, 9, 10, 11, 12^. But whether co-stimulation blockade-induced transplant tolerance and pregnancy share similar transcriptional responses to achieve tolerance to the same antigen remains unsolved.

In pregnancy, tolerance to the semi-allogenic fetus allows for successful parturition as well as repeated pregnancies, whereas in transplantation, exposure to the same alloantigen leads to immunological memory and accelerated rejection of a subsequent graft sharing the sensitizing antigen^13^. Indeed, we reported that prior sensitization to a single antigen expressed by the allograft was sufficient to destabilize transplantation tolerance^13^. The divergent immunological responses to allogeneic conflict prompted us to ask if reencounter of alloantigens in pregnancy after sensitization by paternal skin graft rejection would elicit a recall immune response, or if pregnancy could dominantly constrain memory T cells. We reported that pregnancy was indeed able to restrain memory CD8^+^ T cells toward states of exhaustion^10^. However, the transcriptional changes in antigen-specific naive and memory CD4^+^ Foxp3^neg^ T conventional cells (Tconvs) and Foxp3^pos^ T regulatory cells (Tregs) induced by transplantation and/or pregnancy have not been delineated.

Prior studies probing CD8^+^ and CD4^+^ T cell biology in pregnancy and transplantation have utilized bulk omics sequencing or TCR-transgenic T cells models, but these approaches are challenged by the innate heterogeneity driving T cell biology^14^. Single-cell RNA-sequencing (ScRNA-seq) approaches are beginning to elucidate novel populations of differentiating total or monoclonal TCR-transgenic CD4^+^ T cells^15,16^ but investigations of endogenous polyclonal antigen-specific CD4^+^ T cell populations at the single cell resolution remain technically challenging due to their low frequencies. In this study, we applied ScRNA-seq to profile the transcriptomes of endogenous 2W:I-A^b^ tetramer-binding CD4^+^ T cells that encounter the same 2W:I-A^b^ complex (2W peptide :EAWGALANWAVDSA presented by mouse MHC class II I-A^b^) in settings of semi-allogeneic heart transplantation and pregnancy^17^. Our studies reveal considerable heterogeneity in both Foxp3^neg^ conventional (Tconv) and Foxp3^+^ regulatory (Treg) T cells across models of acute rejection (Rej), anti-CD154-mediated transplant tolerance (Tol), naïve pregnancy at parturition (Pg), and pregnancy after rejection (sensitized pregnancy - SPg). We additionally show that these transcriptional rejection and tolerance responses are conserved in human CD4^+^ T cells. Resolution of these ScRNA-seq signatures in antigen-specific CD4^+^ T cells lays the groundwork for a more precise assessment of their contribution to rejection and tolerance.

## RESULTS

### Donor-reactive CD4^+^ T cells in transplantation rejection and tolerance reveal propensity for divergent effector and regulatory signatures

C57BL/6 (B/6) mice received F1 (BALB/c x B/6) cardiac allografts expressing the model antigen, 2W-OVA (2W.F1)^18^. Naïve tolerant (Tol) allograft recipients were generated by the administration of donor splenocyte transfusion and anti-CD154 on the day of transplantation, followed by 2 additional anti-CD154 doses on post-operative days (POD) 7 and 14. Acute rejection (Rej) recipients received no treatment. Both groups of mice were sacrificed on POD30 and 2W-reactive CD4^+^ T cells were identified with p2W:I-A^b^ tetramers and isolated by flow cytometry for ScRNA sequencing (**Fig. 1A**). We recovered approximately tenfold more 2W^+^ CD4^+^ T cells from pooled Rej mice (n=2) compared to pooled Tol mice (n=6) (**Fig. S1A**).

**Figure 1:**
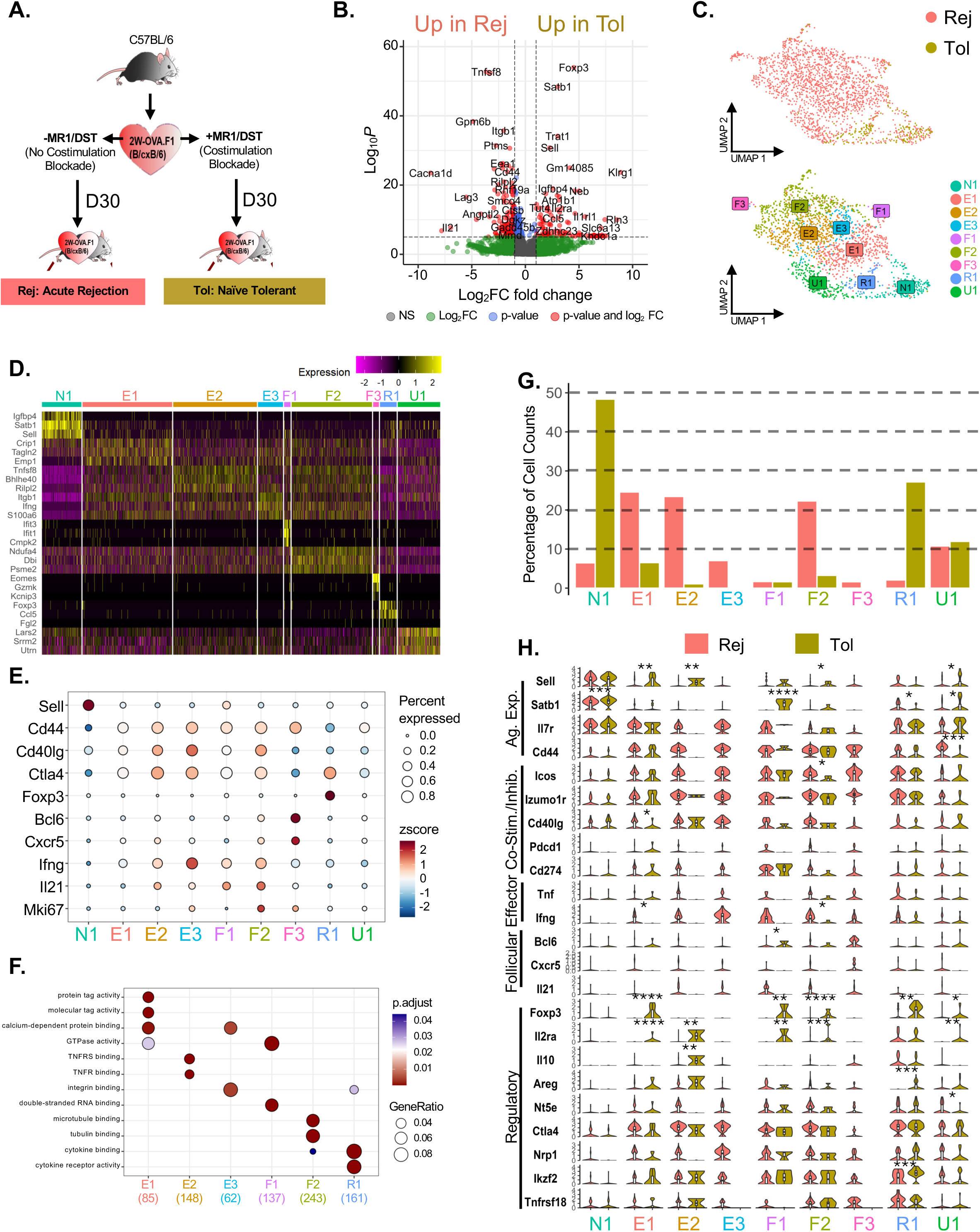
2W-OVA:I-A^b^ CD4^+^ T Cells from Naïve Tolerant heart transplant recipients are enriched for *Foxp3*^+^ while Acute Rejecting 2W-OVA:I-Ab CD4^+^ T Cells are enriched for effector follicular and non-follicular populations. **A.** Experimental design of 2W-OVA-F1 heterotopic allograft transplanted into Acute Rejecting (no immunosuppression; Rej) or Naive Tolerant (with MR1/DST at D0, D7, and D14; Tol) recipients and sacrificed at post-operative D30. **B.** Pseudobulk volcano plot of differentially expressed genes (DEGs) by 2W-OVA:I-A^b^ CD4^+^ T Cells from Rej and Tol. **C.** Dimensionality reduction UMAP split by biological groups, Rej and Tol (top), or nine unbiased clusters (bottom). N1, naive 1; E1, effector 1; E2, effector 2; E3, effector 3; F1, follicular 1; F2, follicular 2; F3, follicular 3; R1, regulatory 1; U1, unclassified 1. **D.** Top n=3 upregulated DEGs in each cluster. Color represents strength of expression. **E.** Dot plot of select canonical markers for CD4^+^ T cell subsets. Size represents percent expression and color represents strength of expression. **F.** Overrepresentation analysis of Gene Ontology terms by cluster. **G.** Bar plot of proportion of unbiased clusters in each experimental group. **H.** Violin plot of canonical marker genes expressed by clusters identified by biological group. Markers are categorized as antigen experience, co-stimulatory/co-inhibitory, effector, follicular, and regulatory. Wilcoxon signed-rank test with Holm correction; *p < 0.05, **p < 0.01, ***p < 0.001, ****p<0.0001.

We first performed pseudobulk analysis of the two transplant groups to reveal major differences between tolerance and rejection. This analysis revealed that *Satb1, Sell, Foxp3, Il2ra,* and *Klrg1* were significantly upregulated in Tol consistent with a quiescent CD4^+^ phenotype and Tregs as previously reported ^13,19^ (**Fig. 1B**). As expected, Rej upregulated *Cd44* as well as *Tnfsf8 (*CD30*), Lag3,* and *Itgb1(*Integrin beta 1) indicating antigen experience and activation. To assess heterogeneity across CD4^+^ T cell populations, we applied UMAP dimensionality reduction and unbiased clustering, which generated nine distinct clusters across both conditions (**Fig. 1C**).

Using the top upregulated and downregulated differentially expressed genes (DEGs) (**Fig. 1D & Fig. S1B**) as well as a select group of markers, we annotated the clusters broadly as: lack of antigen experience (N), effector (E), follicular (F), regulatory (R), or unclassified (U) (**Fig. 1E**). Cluster N1 expressed the highest amount of *Sell* and *Satb1* while Cluster E1 represented a more quiescent effector memory population, marked by upregulation of *Crip1* but minimally expression of canonical effector cytokines, *Ifng or Tnf*. Cluster E2 showed a more activated phenotype, with top differentially expressed genes (DEGs) including *Tnfrsf8*, *Cd40lg*, *Ctla4*, and *Icos*, while Cluster E3 exhibited the most effector-like profile, with the highest *Ifng* and *Cd40lg* expression. Within the follicular category, clusters F1 and F2 co-expressed *Ifng* and *Il21*, whereas Cluster F3 had the highest expression of *Bcl6* and *Cxcr5*. The regulatory R1 cluster had the highest *Foxp3* expression and high *Ctla4* expression. Lastly, cluster U1 was proportionally similar in both Rej and Tol, and was enriched for splicing-associated genes such as *Lars2, Srrm2*, and *Utrn*, suggesting it may represent a transitional cluster (**Fig 1D, G**).

Next, we investigated whether the function of these clusters could be further differentiated based on T cell receptor components (*Cd3e*, Cd3g, *Cd247*, *Cd4*, *Trac* and *Trbc2)*, tyrosine kinases (*Lck* and *Zap70*), or *Thy1* modulation (**Fig. S1C**). Overall, cluster E2 expressed the highest amount of *Trac*, cluster E3 exhibited higher expression of *Cd4* and *Thy1*, while F2 and F3 exhibited higher *CD3g* and *CD3e* respectively, compared to all other clusters. F2 also expressed the highest expression of the tyrosine kinases, *Lck* and *Zap70*. Importantly, these T cell receptor components were expressed at lower levels in clusters N1, R1, and U1 suggesting modular increases in expression of TCR signaling components with the acquisition of effector function in Tconvs.

Given the variable activation markers, T cell receptor components, and signaling cascade expression in each cluster, we next asked if the effector clusters would be overrepresented for biological response, molecular functions, or cellular components from the Gene Ontology database (**Fig. 1F**). We also extended our functional enrichment analysis to interrogate pathways using the Reactome databases of “Metabolism” and “Immune System” (**Figs. S1D-E**). We found E1 was enriched for molecular and protein tag activity, TAK1 complex, and nucleotide and purine salvage metabolic pathways. This was consistent with E1 being a recently activated effector population consistent with its lowest expression of TCR component genes relative to E2 and E3 clusters. E2 was overrepresented by TNFR/TNFRS binding and enriched for TNF and CTLA4 signaling as well as histidine catabolism while E3 was overrepresented by integrin binding, AP-1 activation and glutamate/glutamine metabolism. Thus, E2 and E3 appear to be distinct effectors polarized towards different signaling mechanisms.

F1 was overrepresented by GTPase, double stranded RNA binding and enriched for antiviral mechanisms stimulated by IFN-stimulated genes, in line with high expression of *Ifit1* and *Ifit3,* and for nucleobase and nicotinamide metabolism pathways. F2 was enriched for tubule binding, TCR signaling, and ATP metabolic pathways consistent with ongoing effector responses while F3 was enriched for MHC Class II presentation, NFAT activation, and BH4/cofactor metabolism suggestive of an early Tfh response. Consistent with this, F3 had higher *Bcl6* and *Cxcr5* expression than all other clusters.

The *Foxp3^+^* R1 was overrepresented for cytokine and cytokine receptor activity, while enriching for IL7 and SHC signaling and serine/phosphatidic acid (PA) metabolism. The unclassified cluster U1 was enriched for lipid/phospholipid metabolism pathways, also enriched to a lesser extent in R1. Lipid metabolism has emerged as a crucial regulator of CD4^+^ T Cells differentiation and suggests U1 being a cell transitioning cluster^20^.

We next applied pseudotime trajectory analysis to more systematically distinguish the 9 clusters categories (**Fig. S1F**). Despite proximity in UMAP space, E2/E3 and F2/F3 resolved along distinct pseudotime trajectory branches, with the follicular clusters F2/F3 occupying the most advanced positions along pseudotime indicating that they were most terminally differentiated or most transcriptionally active.

Finally, we demultiplexed the biological groups from each cluster to show that Tol was dominated by antigen-inexperienced cluster N1 and regulatory cluster R1, whereas Rej was dominated by the effector (E1-E3) and follicular (F1-3) clusters (**Fig. 1G**). Notably, Clusters E3 and F3 were restricted to Rej supporting the notion that allograft rejection is mediated by donor-specific T cell and antibody responses. To further interrogate intra-cluster variability of gene expression, we assessed multiple markers broadly classified as antigen experience, co-stimulatory/co-inhibitory and regulatory (**Fig. 1H**). We made the unexpected observation that Clusters E1, F1, and F2 present in Tol expressed *Foxp3*, suggesting a unique propensity towards regulatory cells while also sharing effector and follicular transcriptomes of the E1, F1 and F2 clusters from Rej. A further assessment of genes encoding transcription factors, activation, co-stimulatory, and co-inhibitory markers further advanced our conclusion that E1-3 clusters from Rej were more activated than those from Tol (**Fig. S1G**), and exhibited higher expression of activation markers such as *Tnfrsf4*, *Cd9*, and *Cd200.* Similarly, the follicular F1 and F2 clusters from Rej overexpressed *Gata3* and *Cd28* compared to Tol. Conversely, while Cluster R1 was overall phenotypically similar from both Tol and Rej groups, *Foxp3* and *Areg* expression was significantly higher in Tol compared to Rej.

### Pregnancy expands transcriptionally quiescent fetus-specific Tconvs and Tregs

To test whether pregnancy-induced T cell tolerance imparts the same transcriptome as transplant tolerance, we investigated donor-specific CD4^+^ T cells from post-partum mice. We crossed B/6 naïve virgin (NV) females with 2W-OVA BALB/c males. Spleens and lymph nodes were harvested on post-partum day 0–6 (Pg; n=4) or from NV controls (n=8), and pooled 2W:I-A^b^ CD4^+^ T cells were subject to ScRNA-seq (**Fig. 2A**). Pseudobulk analysis revealed upregulation of *Cd44*, *Ctla4*, and *Il7r* in the Pg group, consistent with antigen experience (**Fig. 2B**). UMAP projection and unbiased clustering revealed considerable transcriptional overlap between groups, with 4 naive-like (N1-4), 2 effector (E1 and E2), 1 Treg (R1) and 1 Unclassified (U1) cluster (**Fig. 2C**). Cluster frequency analysis showed an overall increase in the frequency of E2, R1 and U1 clusters in Pg compared to NV (**Fig. S2A**).

**Figure 2:**
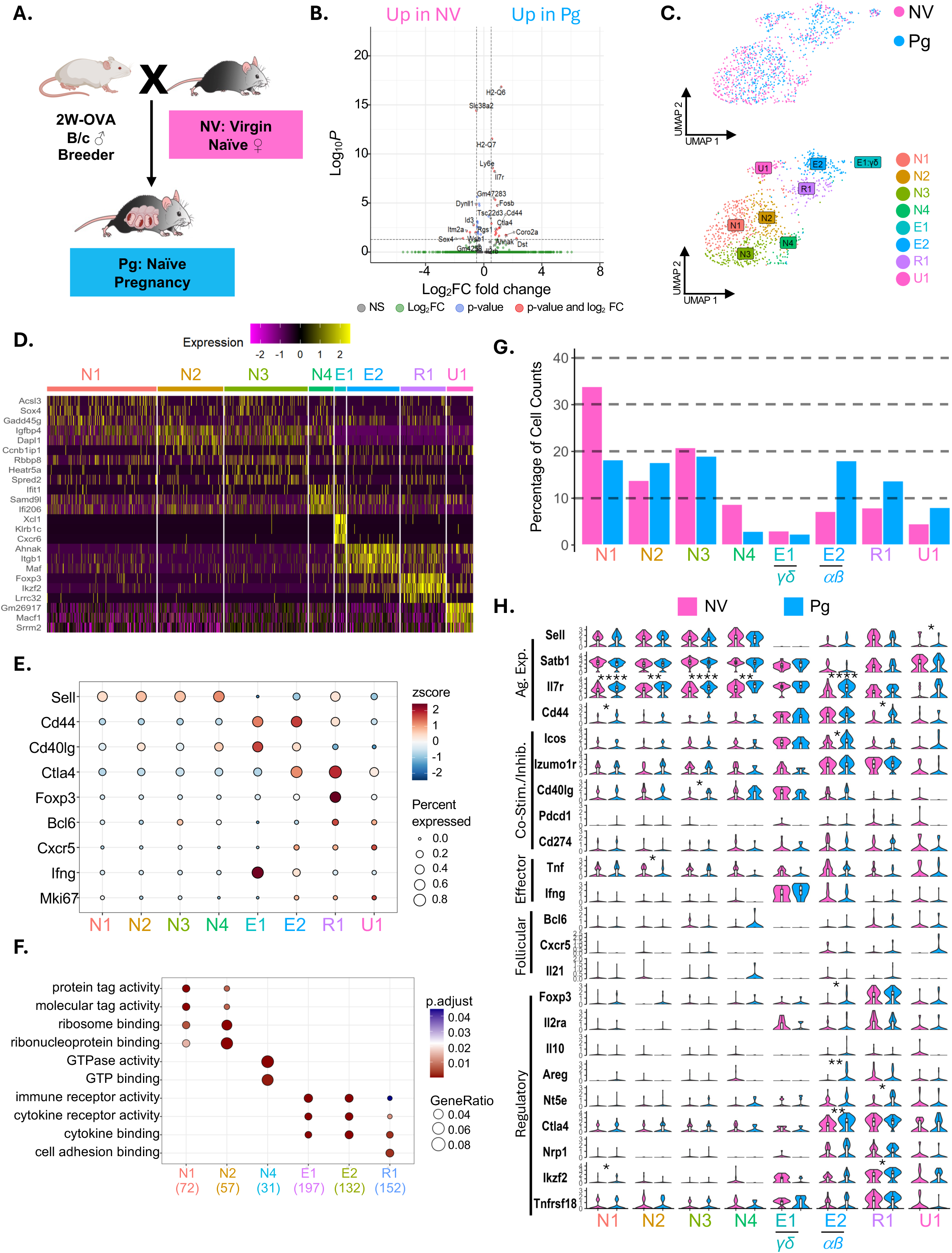
2W-OVA:I-A^b^ CD4^+^ T Cells in Naïve Pregnancy are enriched for an effector and a *Foxp3*^+^ cluster. **A.** 2W-OVA BALB/c male breeders were crossed with Naïve virgin (NV) C57BL/6 females to generate post-partum (Pg) mice that were sacrificed on post-partum D0-6. **B.** Pseudobulk volcano plot of differentially expressed gene by 2W-OVA:I-A^b^ CD4^+^ T Cells from NV and Pg. **C.** Dimensionality reduction UMAP with indicated biological group (top) and eight unbiased clusters (bottom). N1, naive 1; N2, naive 2; N3, naive 3; N4, naive 4; E1, effector gd 1; E2, effector ab 2; R1, regulatory 1; U1, unclassified 1. **D.** Top n=3 upregulated DEGs in each cluster. Color represents strength of expression. **E.** Dot plot of select canonical markers for CD4^+^ T cell subsets. Size represents percent expression and color represents strength of expression. **F.** Over-representation analysis of Gene Ontology terms by cluster. **G.** Bar plot of proportion of unbiased clusters in each experimental group. **H.** Violin plot of canonical marker genes expressed by clusters identified by biological group. Markers are categorized as antigen experience, co-stimulatory/co-inhibitory, effector, follicular, and regulatory. Wilcoxon signed-rank test with Holm correction; *p < 0.05, **p < 0.01, ***p < 0.001, ****p<0.0001.

Cluster annotations based on top upregulated and downregulated DEGs revealed that all four N1-N4 clusters expressed *Sell* and *Satb1* indicating antigen inexperience. N1 was enriched for *Sox4* ^21^, N2 for the naïve T cell marker *Igfbp4* ^22^, and N3 for the negative regulator *Spred2* ^23^, which are consistent with naïve-like phenotypes (**Fig. 2D**, **Fig. S2B**). N4 had *Ifit1* (Interferon-induced protein with tetratricopeptide repeats 1) as a top upregulated gene, but *Sell* expression was high and *Cd44* expression low, consistent with an IFN-conditioned naïve cluster^24, 25^ (**Fig. 2E**). The N1 and N4 clusters were proportionally reduced in Pg.

Cluster U1 lacked *Cd44* expression but expressed the splicing gene *Srrm2* ^26^, suggesting a splicing cluster with no differentiated phenotype. E1 was a relatively minor cluster in both NV and Pg comprising γδ T cells (gamma chain expression not shown) that expressed high *Ifng* and *Cd40lg* (**Fig. 2E-G**) and the highest expression of *Thy1* and *Cd3e* (**Fig. S2C**). The αβ^+^ E2 cluster was expanded in pregnancy and had the highest expression of *Cd44 and Itgb1* (VLA4ß), highly expressed *Cd4* and the tyrosine kinases, *Lck* and *Zap70.* However, E2 minimally expressed *Tnf* and *Ifng*, similar to the E1 cluster in transplant tolerance. R1 had *Foxp3*, *Ikzf2*, and *Lrrc32* as top upregulated DEGs consistent with a regulatory cluster.

Next, we probed Gene Ontology (**Fig. 2F-G**) and the Reactome “Metabolism” and “Immune System” to further understand the E2 and R1 clusters that were proportionally increased in naïve pregnancy (**Fig. S2D-E**). E2 was overrepresented by immune and cytokine receptor activity and enriched for Hyaluronan metabolism and CD28 costimulation. Interestingly, E2 was enriched for the Immunoregulatory interactions between lymphoid and non-lymphoid cells suggesting that E2 might be regulated by cell extrinsic signals, in addition to activated regulatory T cells. The R1 cluster was overrepresented in cytokine and cell adhesion binding, and metabolism that was enriched for PIPs and mitochondrial uncoupling. Consistent with an activated and motile immune response, R1 was also enriched for TNF, NFkB, and IL15 signaling, suggesting a regulatory cluster that was receiving signals and poised for regulatory function. Finally, the U1 cluster from pregnancy was similar to transplantation in the enrichment for phospholipid metabolism but was distinctly enriched for glycosaminoglycan metabolism. In line with findings from the functional gene set analysis, E1, E2, and R1 were furthest along pseudotime, thus iterating their activated phenotype markers (**Fig. S2F**).

Assessment of whether there was significant intra-cluster heterogeneity by biological groups revealed no major transcriptional differences in the N1-N4 clusters from Pg versus NV (**Fig. 2H**). Additionally, E2 clusters from Pg and NV were similar, with those from Pg having increased expression of *Foxp3, Areg and Ctla4.* The R1 cluster was expanded in Pg but was overall transcriptionally similar to R1 from NV, with elevated expression of *Cd44*, *Nt5e*, and *Ikzf2.* Thus, ScRNA-seq analysis showed that while pregnancy expanded the E2 and R1 clusters, they remained remarkably similar to the same clusters from NV. This contrasts with the extensive transcriptional responses in CD4^+^ T cells to the same antigens in the settings of transplant tolerance or rejection.

### Pregnancy after sensitization imparts transcriptional traits of rejection and tolerance in antigen-specific Tconvs and Tregs

The contrasting transcriptional responses in transplantation and naive pregnancy prompted us to examine the transcriptional response of antigen-specific CD4^+^ T cells to pregnancy in dams presensitized by the rejection of paternal skin grafts. To this end, B/6 virgin females were first sensitized by transplantation with 2W-OVA BALB/c skin. At 30–60 days after skin rejection, the mice were mated with 2W-OVA BALB/c males and these sensitized post-partum (SPg; n=4) mice were sacrificed on post-partum days 0–6 along with skin sensitized virgin controls (S; n=3) (**Fig. 3A**). Pseudobulk analysis revealed widespread upregulation of activation markers, TNF family genes, and cytokines in the SPg group while *Il7r* and *Satb1* were most prominent in the S group, consistent with enrichment of resting memory CD4^+^ T cells (**Fig. 3B**).

**Figure 3:**
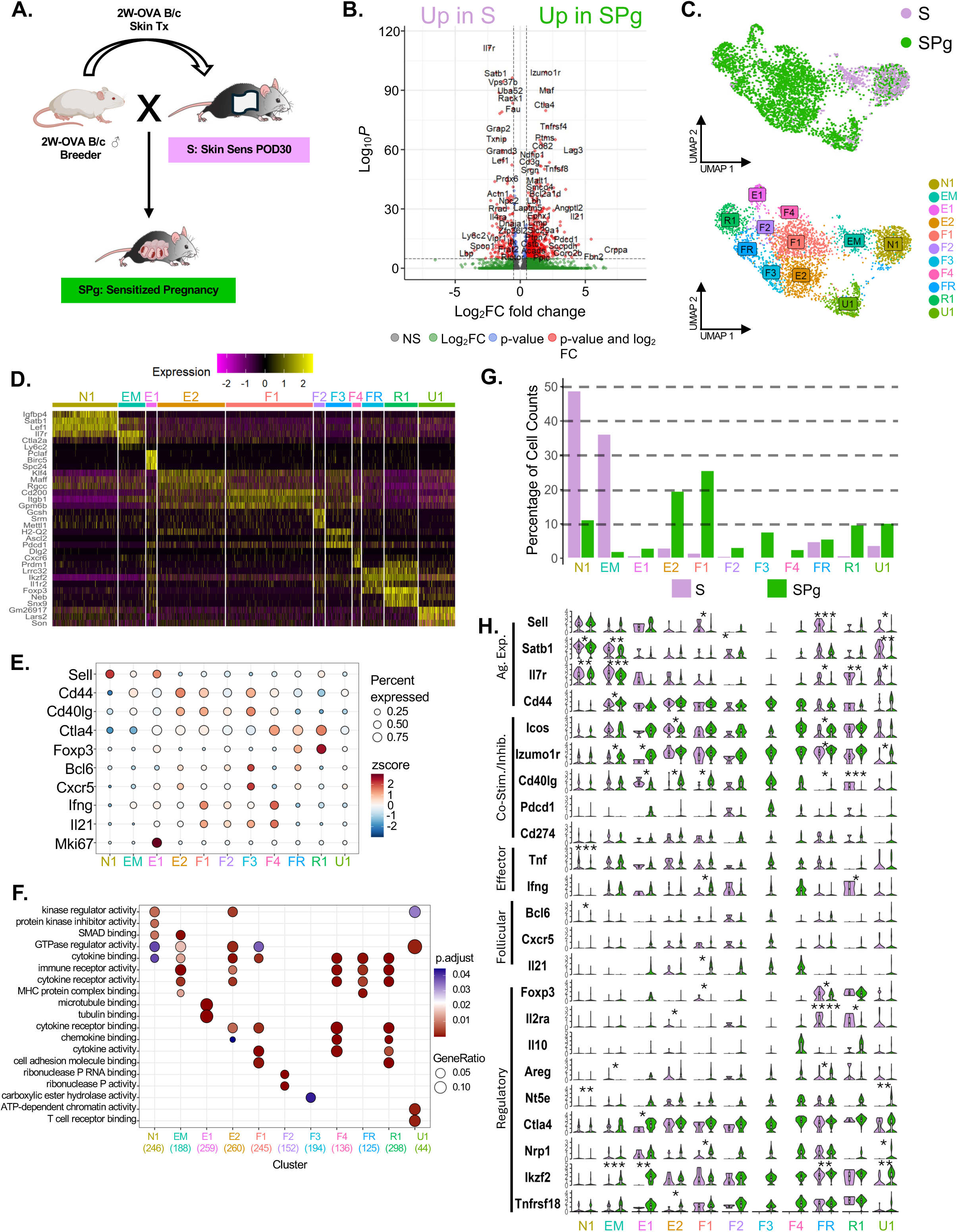
Skin Sensitized Post-Partum mice expand 2W-OVA:I-A^b^ CD4^+^ T Cells into heterogenous effector and regulatory phenotypes. **A.** 2W-OVA BALB/c male breeders were mated with 2W-OVA skin sensitized mice (S) at post-operative D30-60 to generate Sensitized post-partum (SPg) mice, which were sacrificed on post-partum day 0-6. **B.** Pseudobulk volcano plot of DEGs by 2W-OVA:I-A^b^ CD4^+^ T Cells from S and SPg. **C.** Dimensionality reduction UMAP with indicated biological group (top) and 11 unbiased clusters (bottom). N1, naive 1; EM, effector memory; E1, effector 1; E2, effector cluster 2; F1, follicular 1; F2, follicular 2; F3, follicular 3; F4, follicular 4; FR, follicular regulatory; R1, regulatory 1; U1, unclassified 1. **D.** Top n=3 upregulated DEGs in each cluster. Color represents strength of expression. **E.** Dot plot of select canonical markers for CD4^+^ T cell subsets. Size represents percent expression and color represents strength of expression. **F.** Overrepresentation analysis of Gene Ontology terms by cluster. **G.** Bar-plot of proportion of unbiased clusters in each experimental group. **H.** Violin plot of canonical marker genes expressed by clusters identified by biological group. Markers are categorized as antigen experience, co-stimulatory/co-inhibitory, effector, follicular, and regulatory. Wilcoxon signed-rank test with Holm correction; *p < 0.05, **p < 0.01, ***p < 0.001, ****p<0.0001.

Dimensionality reduction and clustering revealed heterogeneity in the CD4^+^ cells, with distinct UMAP regions dominated by S or SPg (**Fig. 3C**). Unbiased clustering identified 11 clusters including an antigen-inexperienced N1 cluster (positive for *Sell, Satb1, Lef1, Igfbp, Cd44^neg^)* and a quiescent memory EM cluster (*Sell^low^, Cd44^pos^, Il7r^+^*, *Ly6c2^+^*) (**Fig. 3D-E & Fig. S3B**). E1 expressed *Mki67* indicating a proliferating cluster, while the more dominant E2 expressed the most *CD44*, as well as the transcription factors, *Klf4* and *Maff* but minimal *Tnf* and *Ifng*, suggestive of an early Th17-like cluster not observed in rejection or naive pregnancy ^27^.

Four follicular-like F1-F4 clusters were identified that expressed varying levels of *Il21*, suggesting that SPg permits the differentiation of Tfh cells. The dominant F1 cluster expressed inhibitory *Cd200* as one of its top DEGs, and co-expressed low levels *Tnf*, *Ifng* and *Il21*. The second most dominant F3 cluster expressed the highest levels of the canonical Tfh genes, *Bcl6*, *Cxcr5, and Pdcd1* while F4 expressed the highest amount of *Il21*, *Ifng, and Ctla4.* Two *Foxp3^+^*Treg clusters were identified: a T follicular regulatory (FR) cluster that expressed *Foxp3, Bcl6,* and *Cxcr5* and a canonical Treg (R1) cluster that highly co-expressed *Foxp3* and *Ctla4.* Finally, an unclassified (U1) cluster was identified that lacked most canonical gene expression but was enriched for the splicing-associated gene *Son* ^28^. Similar to transplant responses, effector clusters E2, F1, F3 and F4 expressed the highest levels of T cell receptor genes (*Cd3e*,*Cd3g*, *Cd4*, *Trac*, and *Trbc2)* and signaling cascade genes *Lck* and *Zap70* (**Fig. S3C**).

Overrepresentation analysis of Gene Ontology terms (**Fig. 3F**) showed considerable overlap between clusters, whereas enrichment analysis against Reactome “Metabolism” and “Immune System” pathways (**Figs. S3D-E**) showed divergent signatures across clusters. E1 was overrepresented by microtubule/tubulin binding and enriched for inflammasomes and nucleotide metabolism and *Mki67* expression indicating a proliferative cluster. In line with *Cd200* expression by E2, GTPase regulator activity and cytokine binding were overrepresented, and IL6 pathways and hyaluronan metabolism were enriched. F1 was overrepresented by cytokine and cell adhesion activity and enriched for keratan sulfate biosynthesis and PD1 signaling; F2 was enriched for noncanonical NFkB signaling and ribonuclease P activity. F3 was enriched for insulin metabolism and AP-1/MAPK pathways while F4 was overrepresented for cytokine and chemokine activity as well as enriched for pyrimidine/histidine catabolism. Regulatory clusters FR and R1 were overrepresented by immune and cytokine receptor activity, but FR lacked pathway enrichment for canonical chemokine and cell adhesion binding pathways suggesting distinct migratory patterns. Furthermore, FR was enriched for ERK inactivation and increased vitamin/cofactor metabolism, while R1 was overrepresented by acyl chain remodeling of phosphatidylinositol (PI) and phosphatidylserine (PS). Finally, the unclassified cluster U1 was overrepresented by ATP-dependent chromatin remodeling activity as well as T cell receptor binding suggesting ongoing differentiation programing. Consistent with our previous models, U1 was also enriched for lipid metabolism suggesting a conserved mechanism of CD4^+^ T cell differentiation.

Pseudotime trajectory analysis revealed that E3 and F1-F4 occupied distinct trajectory branches, consistent with divergent differentiation paths between Tfh and non-Tfh effectors (**Fig. S3F**). To interrogate additional distinctions between follicular clusters F1 and F3 and non-follicular effectors E1 and E2, we enriched these clusters against the Reactome “Immune System” database. Cluster E1 and E2 were enriched inflammasome and IL-6 pathways while F1-F4 were more enriched for immunoregulatory PD-1, immune and NF-kB signaling. Congruent with PD-1^+^ expression by Tfh cells, PD-1 signaling was enriched in the follicular clusters F1 and F3 ^29^. These data underscore the heterogeneity of differentiation pathways taken by Tconvs sensitized by rejection and recalled by pregnancy .

Finally, we interrogated intra-cluster differences between the S and SPg groups. S was dominated by the antigen-inexperienced cluster N1 and quiescent memory cluster EM, while six clusters (E2, F1, F3, F4, R1 and U1) were expanded in SPg (**Fig. 3G**). Antigen-inexperienced cluster N1 had high expression of *Sell* and *Satb1* while EM had higher expression of *Ikzf2* and *Itga4,* with clusters from S and SPg being transcriptionally similar (**Fig. 3H & Fig. S3G**).

Focusing on the clusters that expanded in SPg, E1-SPg had higher *Izkf2, Ctla4, Izumo1r, Tnfrsf4,* and *Casp3* while E2-SPg had higher *Cd40lg* expression compared to E1 and E2 from S, respectively. Consistent with a lack of effector T cell differentiation, neither E1-SPg or E2-SPg increased *Tnf* or *Ifn* compared to S, which was similar to the expanded E2 in Pg mice.

The follicular cluster F1-SPg, compared to F1-S, had higher *Il21, Ifng,* and *Cd40lg* while F1-S had higher *Sell* expression, and F2-SPg had higher *Tnfrsf4* expression. F3 and F4 were restricted to SPg, and had high expression of *Il21, Nt5e, Izumo1r, Icos, Ikzf2* and *Tgfb1.* F3 had higher *Tox2* while F4 had higher *Il10, Il21, Ifng, Itga4, Casp3*, and *Lag3*. Collectively, these transcription profiles are consistent with ongoing Tfh responses in SPg mice at parturition which were not induced in Pg mice Two *Foxp3^+^* Treg clusters were identified in SPg mice. The follicular regulatory (FR) cluster from S versus SPg had distinct activation markers. FR from S had higher *Sell, Bcl2* and *Nr4a1*, while those from SPg had elevated *Icos, Izumo1r* and *Ikzf2*. R1 from S and SPg had similar expression patterns, with R1 from S having higher *Il7r, Cd40lg, Gata3, Il2ra, Ifng*, and *Il7r.* Finally, we noted U1 from SPg having increased expression of *Ikzf2, Izumo1r,* and *Nt5e*, which may point to increased transitioning cells in SPg. Taken together, sensitized pregnancy was associated with marked heterogeneity in expanded Tconvs as well as activated FR and Tregs that was reminiscent of rejection and tolerance, respectively. Notably, this contrasted with the quiescent transcription profiles in Pg mice.

### Conserved effector clusters from mouse models in transplant and pregnancy are enriched in humans

Because the semi-allogeneic graft is rejected but the semi-allogeneic fetus is accepted, we compared effector clusters from transplant (Tx-E2,-E3, -F2) versus pregnancy in naive (Pg) or sensitized (SPg) dams (**Fig. 4A**). Overall, the SPg clusters were most transcriptionally enriched for the three effector Tx clusters, with the two dominant clusters, SPg-E2 and SPg-F1, having the highest module score for the Tx-E2 gene set (**Fig. 4B**). The main Pg expanded non-Treg cluster, Pg-E2, also had a higher module score for Tx-E2 transcriptome. Individually examining the enrichment of SPg-E2 to Tx-E2 revealed a robust positive enrichment score of 0.751 (**Fig. 4C**), whereas Pg-E2 had a negative enrichment score of -0.346. Therefore, SPg-E2 (but not Pg-E2) was similar to Tx-E2 (**Fig. S4A**). Notably, Tx-E3 was the least enriched in both Pg and SPg (**Fig. S4B**), supporting the conclusion that pregnancy does not induce the full differentiation of both naive and memory CD4^+^ T cells to the mature *Ifng*^+^ effectors observed in transplant rejection.

**Figure 4:**
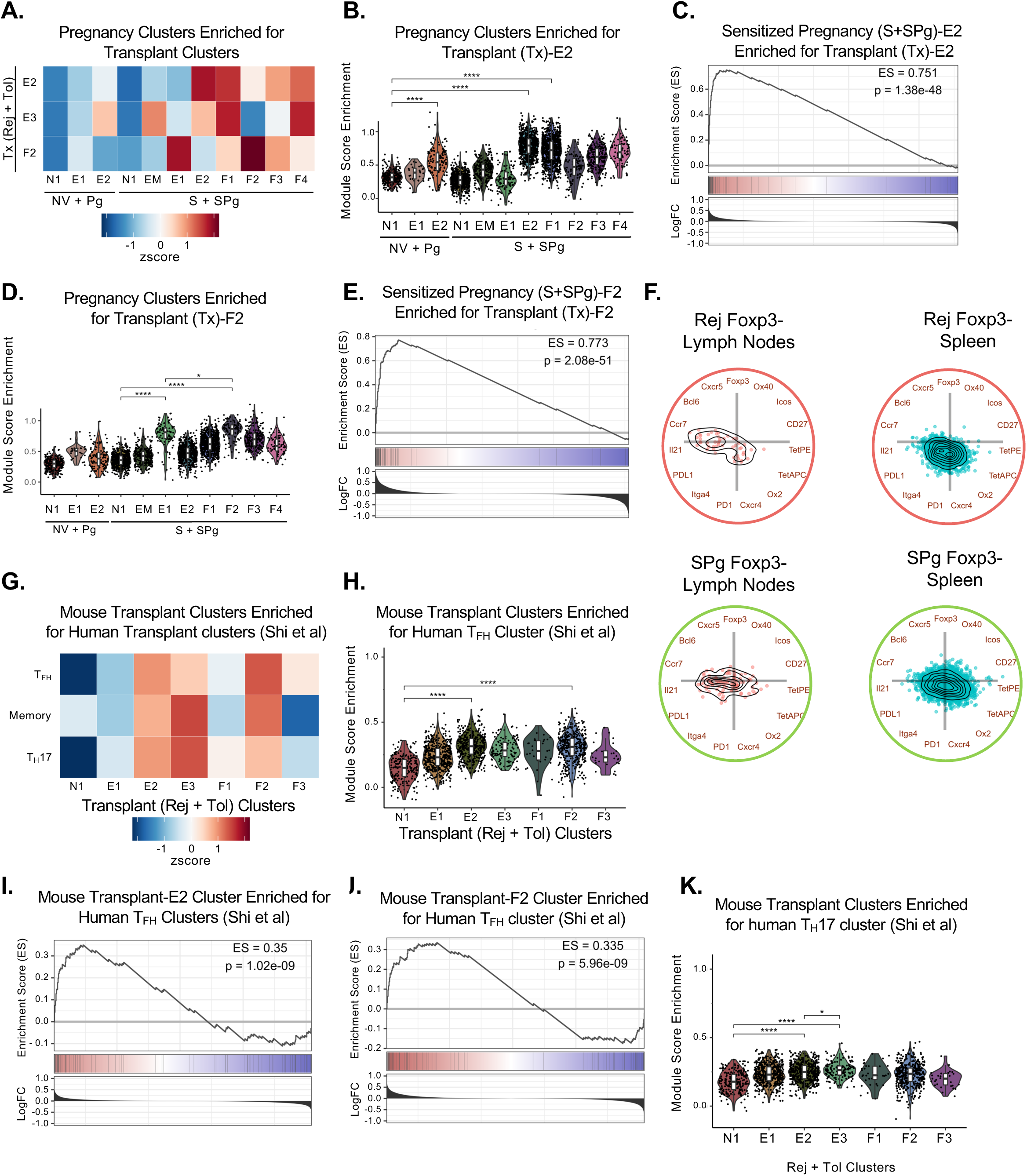
Comparative gene set enrichment analysis of *Foxp3*^−^ 2W-OVA:I-A^b^ CD4^+^ Tconvs in transplant and pregnancy clusters for mouse and human datasets. **A.** Heatmap of scaled enrichment of Naïve pregnancy (from NV & Pg; x-axis) and Sensitized pregnancy (from S & SPg; x-axis) effector clusters against transplant (from Rej + Tol; y-axis) effector clusters. Color represents strength of expression. **B.** Module score of Naïve pregnancy (from NV & Pg) and Sensitized pregnancy (from S & SPg) effector clusters against transplant effector cluster Tx-E2 (from Rej & Tol). Wilcoxon signed-rank test with Holm correction; *p < 0.05, **p < 0.01, ***p < 0.001, ****p<0.0001. **C.** Gene Set Enrichment Analysis (GSEA) of effector E2 cluster from sensitized pregnancy (from S & SPg) versus transplant Tx-E2 (from Rej & Tol). **D.** Module score of Naïve pregnancy (from NV & Pg) and Sensitized pregnancy (from S & SPg) effector clusters against transplant effector cluster Tx-F2 (from Rej & Tol). Wilcoxon signed-rank test with Holm correction; *p < 0.05, **p < 0.01, ***p < 0.001, ****p<0.0001. **E.** GSEA of follicular F2 cluster from sensitized pregnancy (from S & SPg) versus transplant F2 (from Rej & Tol). **F.** Radviz of *Foxp3*-2W-OVA:I-A^b^ CD4^+^ from the lymph nodes or spleens of Rej and SPg recipients, showing flow cytometry protein expression for follicular, effector, and 2W-OVA tetramer markers (PE and APC). **G.** Heatmap of scaled enrichment of transplant clusters (from Rej & Tol; x-axis) against *Shi et al.* ^31^ graft-infiltrating clusters of kidney transplant rejection patients (y-axis). **H.** Module score enrichment of transplant clusters (from Rej & Tol) enriched for *Shi et al.* ^31^ “Tfh” CD4^+^ gene set. Wilcoxon signed-rank test with Holm correction; *p < 0.05, **p < 0.01, ***p < 0.001, ****p<0.0001. **I.** GSEA of Tx-E2 (from Rej & Tol) enriched for *Shi et al.* ^31^ “Tfh” CD4^+^ gene set. **J.** GSEA of Tx-F2 (from Rej & Tol) enriched for *Shi et al.* ^31^ “Tfh” CD4^+^ gene set. **K.** Module score enrichment of transplant clusters (from Rej & Tol) enriched for *Shi et al.* ^31^ “Th17” CD4^+^ gene set. Wilcoxon signed-rank test with Holm correction; *p < 0.05, **p < 0.01, ***p < 0.001, ****p<0.0001.

Next, we focused on the Tfh-like Tx-F2 cluster which had a positive enrichment with several SPg effector clusters, whereas the sole Pg-E2 cluster was not enriched for Tx-F2. These observations are congruent with pregnancy-induced fetus-specific antibody (FSA) responses in naive mice being a weaker GC-independent response compared to the donor-specific antibody-responses (DSA) in Rej (**Fig. 4D**)^8^. SPg-F2 had the most positive enrichment of 0.773 with Tx-F2, consistent with a robust positive conserved follicular signature (**Fig. 4E**). When SPg non-Tfh clusters were compared to follicular Tx-F2, SPg-E1 was positively enriched (**Fig. S4C**) whereas S-E2 was negatively enriched (**Fig. S4D**). These data confirm that pregnancy in sensitized dams permits broad Tfh differentiation that overlapped with rejection.

Finally, we confirmed that the transcriptional observations reflect protein expression by generating a spectral flow cytometry panel that probed for several T effector and follicular markers. To highlight the most dominant markers and phenotypes at the single cell resolution, we used Radviz visualization whereby markers are arranged along a circumference based on similarity of relative expression, and differentiated cells are positioned towards the most dominant markers and away from the center ^30^. Overall, we found that Rej 2W:I-A^b^ Tconvs from the lymph nodes and spleen were slightly more polarized compared to SPg Tconvs, and that non-Tfh markers more polarized in the spleen (**Fig. 4F**). Additionally, polarization of splenic Tconvs from both Rej and SPg towards the PE and APC tetramers suggest influence by TCR avidity, and were diametrically opposite from the follicular markers, Bcl6 and Il21. Indeed, traditional flow gating confirmed more tetramer MFI high cells were found in the spleen than lymph nodes informing the location of effector T cells with GC follicular T cells enriching in the lymph nodes and non-Tfh effector cells accumulating in the spleen (**Fig. S4E**).

We next tested whether the effector clusters of antigen-specific Tconvs in mice shared transcriptional profiles reported for human transplantation. Using a ScRNA dataset from kidney graft-infiltrating T cells at rejection (KTx) from *Shi et al, 2023* ^31^ we confirm that the mouse transplant (Tx) clusters E2, E3, and F2 were enriched for human kidney transplant rejection Tfh, Memory, and Th17 gene sets (**Fig. 4G**). Tx-E2 and Tx-F2 clusters had the highest module scores for KTx Tfh (**Fig. 4H**) and GSEA of Tx-E2 and Tx-F2 showed positive gene set enrichment scores (ES=0.35 and 0.335 respectively) for KTx Tfh (**Fig. 4I-J**). When compared to KTx Th17, Tx-E2 and Tx-E3 had modestly higher modules scores than the other mouse clusters (**Fig. 4K**), with positive enrichment of Tx-E2 cluster for KTx Th17 (ES=0.215). In contrast, Tx-E3 had a negative enrichment score (ES= -0.186), consistent with a lack of a robust Th17 signature in mouse heart transplant rejection (**Fig. S4F-G**). Compared to the KTx Memory cluster, none of the transplant-induced clusters had significantly enriched module scores consistent with an on-going antibody response (**Fig. S4H**). Finally, we extended our analysis to ask if the mouse pregnancy clusters were enriched for a human dataset from *Moldenhauer et al, 2024* ^32^ of Tconvs from Early Pregnancy Failure. Consistent with successful pregnancies in both our mouse models, they were not enriched (**Fig. S4I**).

### Heterogeneous follicular T cells associated with alloantibody responses in acute rejection and pregnancy after transplant rejection

The conserved follicular signature in SPg mice was consistent with a strong anti-2W-OVA IgG response that was higher than anti-2W-OVA IgG in Rej and S known to affect graft survival^33^ (**Fig. 5A**). To further probe for heterogeneity within the conserved Tfh phenotypes, we performed a sub-analysis of CD4^+^ Tfh-like cells expressing *Bcl6*, *Cxcr5*, *Il21* or *Pdcd1* from Rej and SPg mice (**Fig. 5B**). Nine follicular-like clusters were identified and verified to be “bona fide” follicular T cells based on module scores against a published “Full Tfh” gene set from *Podesta et al, 2023* ^15^. All clusters, except SF and P1, demonstrated high enrichment (**Fig. 5C**). Differential gene expression analysis revealed that SF was enriched for the splicing and apoptosis-associated ^34^ gene *Lars2* suggesting a transitioning state (**Fig. 5D & Fig. S5A**). Enrichment analysis using “Progenitor, Tfh versus Tfr, and Tfr” gene sets from *Podesta et al, 2023* ^15^ and *Huo et al, 2019* ^35^ confirmed P1 as a progenitor-like cluster and pFR as Tfr, with the remaining clusters consistent with conventional Tfh phenotypes (**Fig. S5B**).

**Figure 5:**
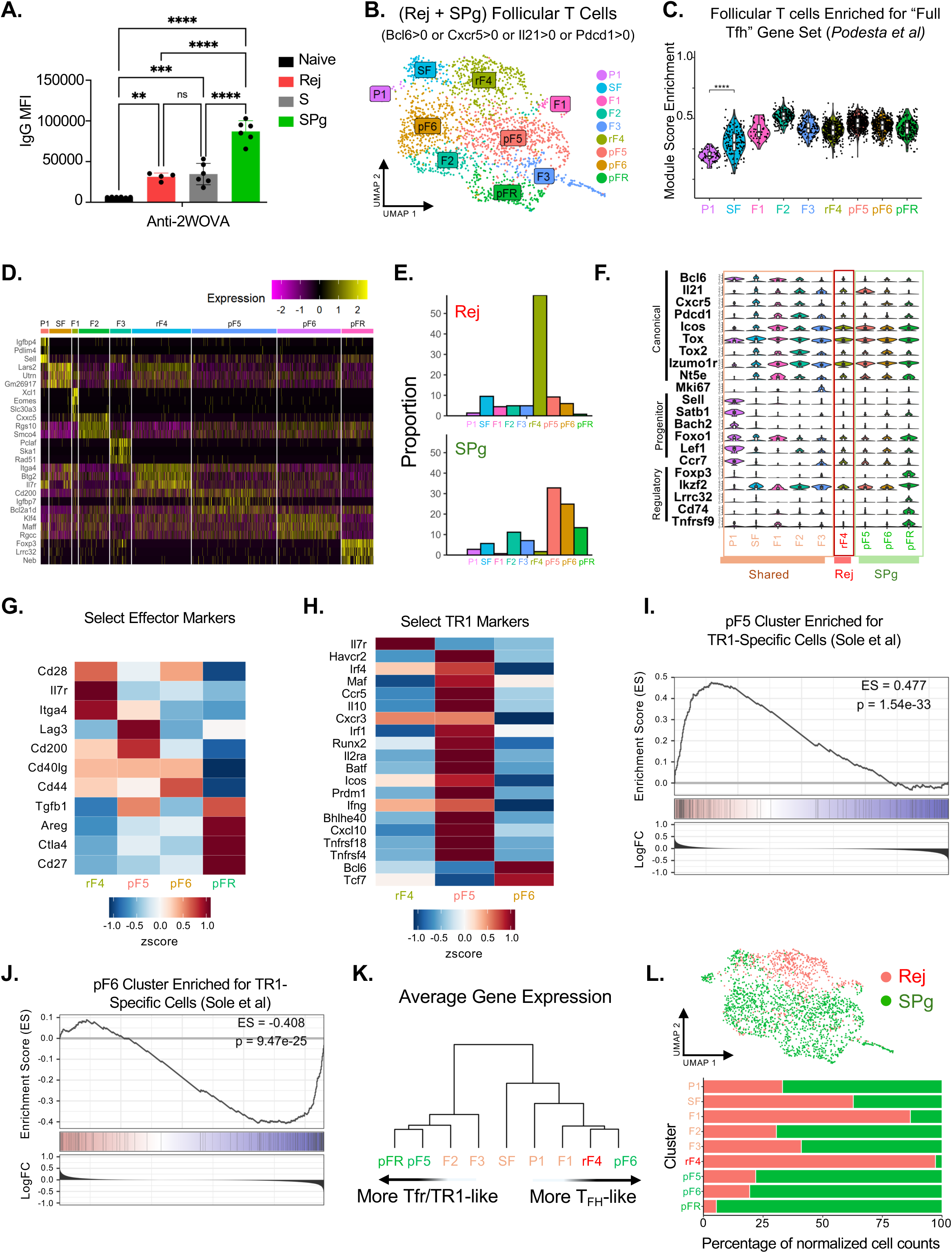
Acute Rejection and Sensitized Pregnancy elicit distinct 2W-OVA:I-A^b^ CD4^+^ Follicular T cell phenotypes. **A.** Mean Fluorescence Intensity (MFI) of anti-2W-OVA IgG in the sera from Naïve, Rej, S, and SPg. Ordinary 1-way ANOVA *p < 0.05, **p < 0.01, ***p < 0.001, ****p<0.0001. **B.** UMAP of 2W-OVA:I-A^b^ CD4^+^ Follicular T Cells by unbiased clusters from Rej and SPg. Cells were subset as: Bcl6>0 or Cxrc5>0 or Il21 >0 or Pdcd1 >0. P1, [Progenitor 1]; SF, [Splicing Follicular]; F1, [Follicular 1]; F2, [Follicular 2]; F3, [Follicular 3]; rF4, [rejecting Follicular 3]; pF5 [pregnancy Follicular 5]; pF6 [pregnancy Follicular 6]; pFR [pregnancy Follicular Regulatory]. **C.** Violin plot of module score by clusters against *Podesta et al.*^15^ “Tfh Full” gene set. Wilcoxon signed-rank test with Holm correction; *p < 0.05, **p < 0.01, ***p < 0.001, ****p<0.0001. **D.** Top n=3 upregulated DEGs by cluster. Color represents strength of expression. **E.** Bar plot of unbiased clustering broken down by biological group. **F.** Violin plot of select Follicular marker genes including canonical, progenitor, and regulatory expressed by each unbiased cluster. **G.** Scaled heatmap of select effector markers for clusters rF4, pF5, pF6, and pFR. Color represents strength of expression. **H.** Scaled heatmap of select TR1-related markers for clusters rF4, pF5, pF6. Color represents strength of expression. **I.** GSEA of pF5 on *Solé et al.*^36^ TR1-Specific gene set. **J.** GSEA of pF6 on *Solé et al.* ^36^ TR1-Specific gene set. **K.** Phylogenetic tree of average gene expression by cluster. **L.** Proportion of cells from biological groups in each cluster.

Cluster proportion analysis revealed a single rejection (rF4) cluster dominant in Rej, and three increased clusters, pF5, pF6, and pFR in SPg (**Fig. 5E**). Apart from P1 expressing more progenitor markers and pFR expressing more regulatory markers, most clusters similarly expressed canonical follicular markers (**Fig. 5F**). The transplant-dominant rF4 expressed high *Il7r* and *Itga4* consistent with a memory-like state as the T cells were analyzed on D30 post-transplantation after allograft rejection on ∼D10-12^13^ (**Fig. 5G**). In contrast, pF5, pF6, and pFR, which were dominant in SPg, were low for *Il7r* consistent with an ongoing anti-2W-OVA IgG response.

We noted that pF5 expressed more *Lag3* and *Tgfb1* compared to pF6 and rF4, which is reminiscent of *Foxp3*^−^ *Il10*^+^ TR1 cells. Indeed, pF5 compared to rF4 and pF6 expressed higher levels of many TR1-associated genes, including *Il10* (**Fig. 5H**). To test this possibility further, we used a TR1-specific gene set from *Solé et al, 2023* ^36^, and observed that pF5 had a higher module enrichment score than rF4 and pF6 (**Fig. S5C**). GSEA of the TR1-specific gene set showed that cluster pF5 and pFR were positively enriched for TR1 genes, with scores of ES=0.477 and 0.458 respectively (**Fig. 5I, Fig. S5D**); in contrast, pF6 and rF4 were negatively enriched (**Fig. 5J, Fig. S5E**). Furthermore, both pF5 and pFR exhibited the highest number of predicted cell-cell interaction strength supporting their regulatory identity (**Fig. S5F**). A direct comparison of the most differentially expressed genes between pF5 and pFR revealed that despite a shared TR1 signature, pF5 highly expressed *Cd30, Ifng, Il21,* and *Cd200* compared to pFR, while pFR was enriched for regulatory genes *Foxp3* and *Ikzf2*. These data reinforce the conclusion of two distinct follicular-like regulatory cell populations in SPg (**Fig S5G**).

A phylogenetic tree based on average gene expression revealed that follicular CD4^+^ T cell heterogeneity could be placed on a divergent spectrum, ultimately leading to Tfh-like or Tfr/TR1-like phenotypes (**Fig. 5K**). Pseudotime and RNA velocity analyses corroborated this divergence, with pregnancy-dominant pF6 and pFR further along pseudotime than transplant-dominant rF4 (**Fig. S5H**). These pF6, pFR and rF4 clusters also diverged in intronic velocity directionality (**Fig. S5I**) . Taken together, these analyses underscore the complexity of the CD4^+^ response in SPg that manifests as heterogeneity in both Tfh and regulatory responses that ensures successful pregnancy despite robust fetus-specific antibody responses (**Fig. 5L**).

### Resolving Foxp3^+^ Treg heterogeneity in transplantation and pregnancy

Expanded clusters of *Foxp3^+^* T cells in all our experimental models prompted a focused analysis on whether they were transcriptionally conserved across pregnancy and co-stimulation blockade-induced transplantation tolerance. We performed a subset analysis by selecting all cells with *Foxp3* expression (**Fig. 6A**). Clustering revealed distinct enrichment of 6 clusters that were annotated based on canonical markers, and heterogenous expression of *Foxp3* (**Fig. 6B-C & Fig. S6A**). Each experimental group harbored at least 4 clusters, with quiescent Central Tregs dominant in Tol and S (at 30-60 days post-sensitization with skin graft). Unexpectedly, despite ten-fold fewer *Foxp3*^+^ cells recovered from Rej, the proportion of Treg clusters in Tol and Rej were similarly dominated by *Foxp3*-high and Effector-high clusters, with an additional expanded Tfr cluster in Rej. In contrast, Pg was dominated by Central Tregs, with the remaining 40% comprising *Foxp3*-high, Effector-high and Tfr clusters. Most remarkable was pregnancy in sensitized dams (SPg), which resulted in the expansion of all 5 non-Central Treg clusters with the Effector-low, Effector-high and Tfr clusters dominating.

**Figure 6:**
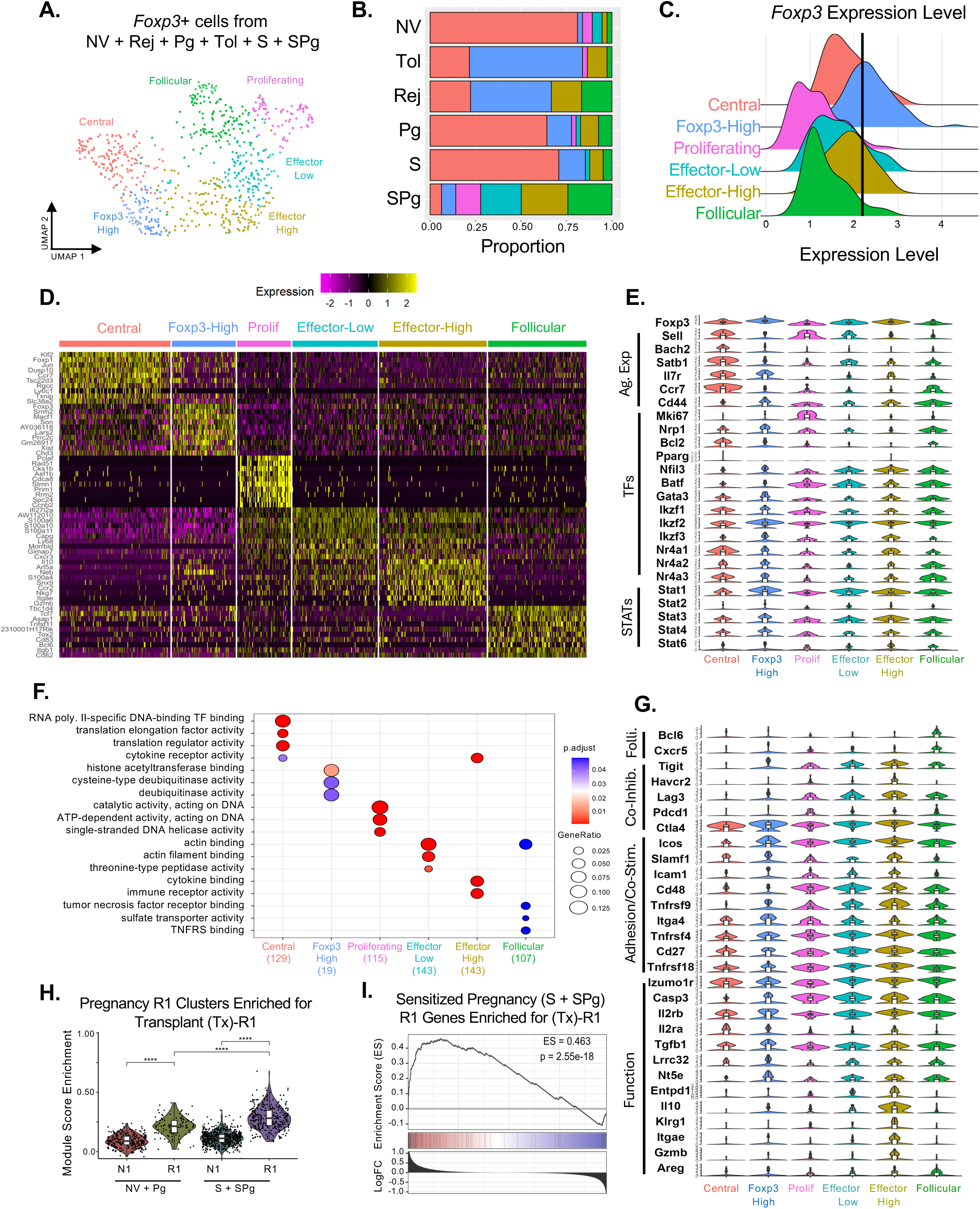
*Foxp3*^+^ 2W-OVA:I-A^b^ CD4^+^ subset analysis across transplantation and pregnancy. **A.** UMAP of 2W-OVA:I-A^b^ CD4^+^ T Cells with reads of *Foxp3*>0 subset from all biological groups were re-clustered and annotated. Colors indicate different types of Foxp3^+^ T cells. **B.** Stacked barplot of *Foxp3*^+^ clusters broken down by biological group. **C.** Ridgeplot of *Foxp3* expression for each annotated cluster. **D.** Heatmap of top n=10 upregulated differentially expressed genes. Color represents strength of expression. **E.** Violin plot by cluster of select Regulatory marker genes including antigen experience, canonical regulatory, transcription factors (TFs), STAT Proteins, and activation/function. **F.** Overrepresentation analysis of Gene Ontology terms by cluster. **G.** Violin plot by cluster of select Regulatory marker genes including antigen experience, canonical regulatory, transcription factors (TFs), STAT Proteins, and activation/function. **H.** Module score of clusters N1 and R1 from naïve pregnancy (NV & Pg) and sensitized pregnancy (S &SPg) N1 and R1 against our transplant *Foxp3*^+^ cluster (Tx-R1). Wilcoxon signed-rank test with Holm correction ; *p < 0.05, **p < 0.01, ***p < 0.001, ****p<0.0001. **I.** GSEA of R1 (from S & SPg) versus Tx-R1 (from Rej & Tol).

Differential gene expression analysis confirmed that Central Tregs were most antigen inexperienced and expressed *Sell* but not *CD44*, while Proliferating Tregs showed strong cell cycle signatures including *Mki67*. The *Foxp3* High cluster was uniquely *Mif* negative/low and enriched for RNA splicing genes. Follicular Tregs expressed *Bcl6* and *Cxcr5,* Effector-low Tregs upregulated S100-family genes while Effector-high Tregs highly expressed genes implicated in Treg function including *Entpd1, Gzmb, Itgae, Il10, Klrg1*, and higher *Tcf7* and *Tnfrsf11* (**Fig. 6D-E & Fig. S6B**). GO enrichment confirmed these annotations (**Fig. 6F**). TCR component genes were most abundant in the Follicular and Effector high/low Treg clusters, mirroring the modular regulation observed in effector Tconvs (**Fig. S6C**). A deeper look at functional, adhesion, co-stimulatory/co-inhibitory, and follicular genes revealed that Effector-High Tregs expressed more functional Treg genes than Effector-Low, Follicular, and Proliferating Tregs, despite sharing expression of many other genes (**Fig. 6G**).

Metabolism enrichment analysis using the Reactome database revealed the Foxp3-high Tregs were enriched for lipid metabolism pathways reminiscent of the Unclassified clusters across our models (**Fig. S6D**). Proliferating and Effector low Tregs shared insulin pathways while Follicular and Effector low Tregs shared a fatty acid pathway enrichment. Effector high Tregs enriched for serine and phosphate bond hydrolysis pathways reinforcing an active effector phenotype. Effector high and Effector low Tregs shared lipid particle organization pathway enrichment. Trajectory analysis suggested that Proliferating Tregs bifurcate into Follicular or Effector-Low lineages, with the latter transitioning into Effector-High cells (**Fig. S6E**). Intronic RNA velocity analysis highlighted *Foxp3*-high Tregs as having the highest pseudotime (**Fig. S6F**) suggesting increased transcriptional priming, while gene-specific RNA velocity plots (e.g., *Itga4* and *Klrg1*) impute an effector-like trajectory (**Fig. S6G**).

To evaluate transcriptional similarity across transplantation (Tx) and pregnancy in naive (Pg) versus sensitized (SPg) mice, we compared their Treg gene sets. Module score and GSEA enrichment against the transplant-derived Tx-R1 cluster revealed that while both Pg-R1 and SPg-R1 showed positive enrichment, SPg-R1 was more enriched (**Fig. 6H–I & Fig. S6H**). Indeed, splitting the Effector High cluster by biological group and examining *Il10* and *Gzmb* expression revealed sensitized pregnancy generates a more robust functional signature than naïve pregnancy but comparable to Tol and Rej^37^ (**Fig. S6I**).

### Conserved transplantation and pregnancy Foxp3^+^ clusters are preferentially enriched in lymphoid organs and human datasets

Tregs function to dampen T cell priming in secondary lymphoid organs in LNs and spleen ^38^. Thus, we compared our dataset with a reference scRNA-seq study from *Miragaia et al, 2019* ^39^ that defined the transcriptomes of lymphoid tissue-specific Treg states in mice. Our antigen-specific Central, Effector-Low, Effector-High, and Follicular Tregs were enriched in the Miragaia splenic Treg gene signatures, while the Proliferating and to a lesser extent, Follicular Tregs, were enriched for Miragaia brachial lymph node effector signatures (**Fig. 7A**). Notably, our Central Tregs had a high positive enrichment score of 0.867 (**Fig. 7B**) and a robust positive module score (**Fig. S7A**), indicative of a conserved resting Treg signature in secondary lymphoid organs. When scored against the Miragaia splenic effector gene set, our Effector-High and -Low Treg clusters showed strong positive module scores while our Proliferating and *Foxp3* High clusters were only modestly enriched (**Fig. 7C**). Finally, our Proliferating cluster had a positive module score and positive enrichment for the Brachial lymph node effector gene set (**Figs. S7B & S7C**). Taken together, these data may provide insights into the location of Treg differentiation and/or accumulation within different secondary lymphoid organs.

**Figure 7:**
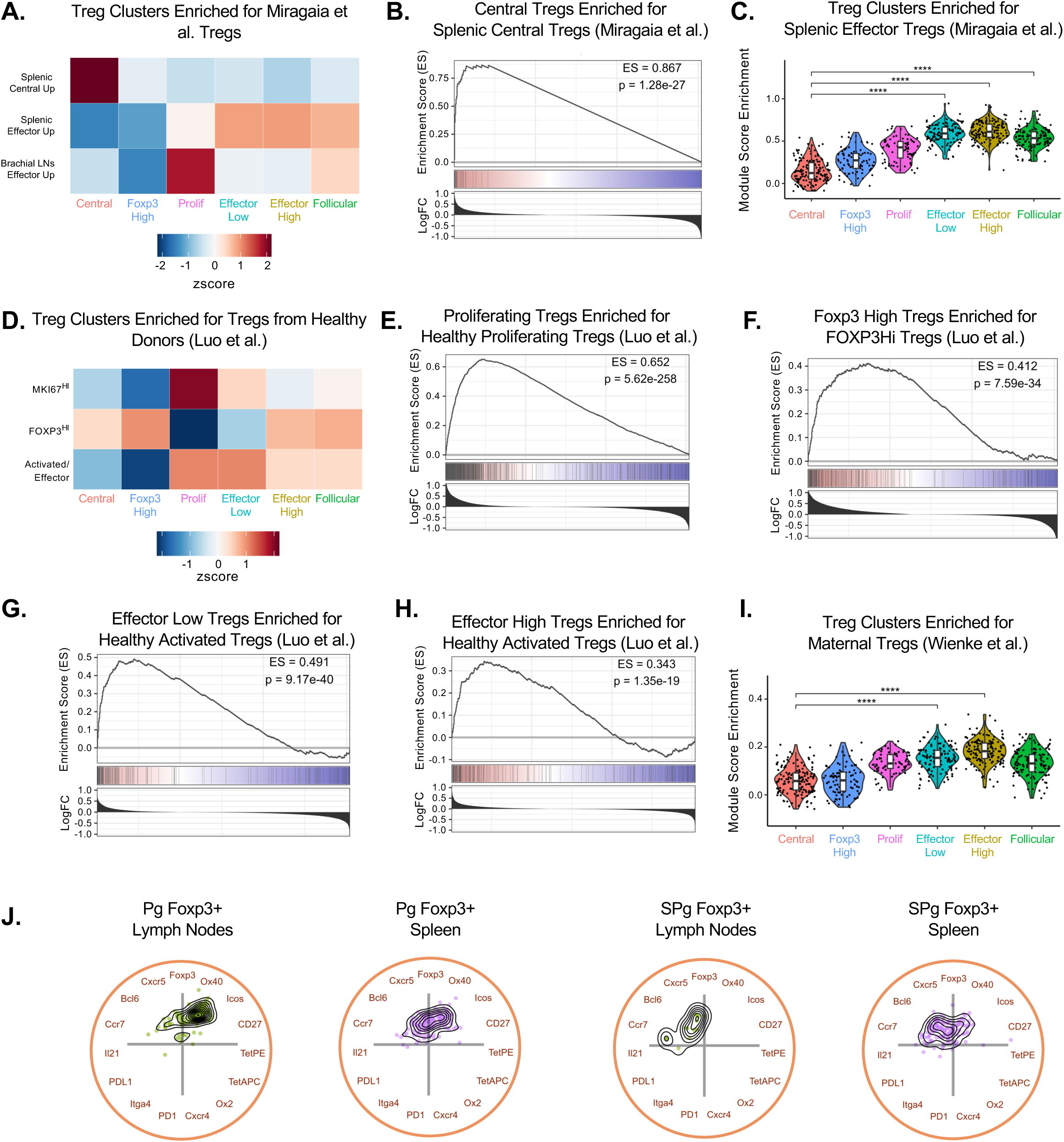
Comparative gene set enrichment analysis of transplant and pregnancy *Foxp3*^+^ 2W-OVA:I-A^b^ CD4^+^ clusters for mouse and human Treg datasets **A.** Heatmap of scaled enrichment by cluster of enrichment for the *Miragaia et al.* ^39^ gene set. Color represents strength of expression. **B.** GSEA of Central Tregs on *Miragaia et al.* ^39^*“*Splenic Central Tregs.” **C.** Module score of Treg clusters against *Miragaia et al.* ^39^ *“*Splenic Effector Tregs.” Wilcoxon signed-rank test with Holm correction ; *p < 0.05, **p < 0.01, ***p < 0.001, ****p<0.0001. **D.** Heatmap of scaled enrichment by cluster of enrichment for the “Healthy Donor Tregs” gene set from *Luo et al.* ^40^. Color represents strength of expression. **E.** GSEA of Proliferating Tregs on *“*MKI67 Hi” gene set from *Luo et al.* ^40^. **F.** GSEA of *Foxp3* High Tregs on*“FOXP3* Hi” gene set from *Luo et al.* ^40^. **G.** GSEA of Effector Low Tregs on *“FOXP3* Activated/Effector” gene set from *Luo et al.* ^40^. **H.** GSEA of Effector High Tregs on *“FOXP3* Activated/Effector” gene set from *Luo et al.* ^40^. **I.** Module score of clusters against *“*Maternal Uterine Tregs” from *Wienke et al.* ^41^. Wilcoxon signed-rank test with Holm correction ; *p < .05, **p < 0.01, ***p < 0.001, ****p<0.0001. **J.** Radviz of Pg and SPg *Foxp3*^+^ Tregs from lymph node or spleen.

We next conducted cross-species validation using a healthy human peripheral blood Treg dataset from *Luo et al, 2021* ^40^, which also revealed conserved signatures. Mouse Proliferating Treg cluster had the highest module score (**Fig S7D**) and was positively enriched for the human MKI67^hi^ population (ES=0.65; **Fig. 7E**), while the mouse *Foxp3* High cluster was positively enriched for the human *FOXP3*^hi^ subset (ES=0.412; **Fig. 7F**). Mouse Effector-Low cluster corresponded closely to the Activated/Effector subset; this is congruent with the human Treg gene set being derived from healthy donors and the Effector-Low cluster being prevalent in naive mice (**Fig. 7G–H**). We also compared mouse Treg clusters to gene sets from peripheral blood Tregs derived from human acute Graft vs Host Disease (GVHD) patients, which were classified as activated or proliferating. We now observed high enrichment and module enrichment scores for mouse Proliferating, Effector and Follicular Tregs for GVHD Activated Tregs, and our Proliferating Tregs were most enriched for the GVHD Proliferating Tregs (**Fig S7E-F**). These observations further confirm conserved signatures of Treg activation and proliferation across species in health and disease states.

To further validate Treg responses in pregnancy, we enriched our clusters against a gene set of unstable Tregs associated with Early Pregnancy Failure ^32^. However, we did not observe module score enrichment in any of our Tregs clusters suggesting our Treg clusters were stable (**Fig. S7G**). Consequently, we probed the transcriptome of healthy, pregnancy-specific Tregs from *Wienke et al,* ^41^ who reported a gene set of activated, maternal uterine Tregs. Effector High cluster had the highest module scores followed by the Effector Low, (**Fig. 7I**) which was recapitulated by modest GSEA enrichment score of 0.26 and 0.182, respectively (**Figs. S7H and S7I**). These data suggest that stable effector Tregs in successful pregnancy are transcriptionally conserved in mice and humans and can be distinguished from unstable human Tregs.

Based on the findings of Wienke et al. ^41^ that *Ox40* was a conserved activation marker of uterine Tregs during pregnancy, we examined OX40 as well as select effector and follicular markers on Tregs from Pg and SPg. From both lymph node and spleen, Radviz visualization of these flow cytometry markers confirmed OX40 and Foxp3 were grouped next to each other indicating co-expression. It also revealed that Pg Tregs were polarized toward an OX40^+^ Icos^+^ phenotype, whereas SPg Tregs displayed dual OX40^+^ and Bcl6^+^/Ccr7^+^ polarization (**Fig. 7J**), consistent with SPg Tregs having the greatest representation of Effector-High and Follicular subsets. These observations support the conclusion that sensitized pregnancy elicits the most functionally diverse and activated Tregs that are similarly reflected in the spleen and lymph node.

## DISCUSSION

Herein, through ScRNA phenotyping and a comparative analysis approach, we demonstrate the heterogeneity of endogenous, antigen-specific CD4^+^ T cell responses that were distinct and overlapping in heart transplantation, pregnancy, and pregnancy after sensitization by transplant rejection. Transplant rejection was characterized by follicular and non-follicular *Ifng*^+^ effectors, whereas transplant tolerance was associated with the expansion of *Foxp3*^+^ Tregs. These follicular and Th1-like effectors were also enriched in a human, kidney-infiltrating gene set of rejecting patients^31^, highlighting the relevance and overlapping phenotype between mice and humans in rejection. That antigen-specific CD4^+^ T effector cell signatures enrich within human *in-situ* gene sets supports the relevance of the effector heterogeneity we have articulated in mice. A surprising observation was that in Tol, the same follicular and non-follicular effector clusters expressed *Foxp3*, while conversely, the Treg cluster from Rej had lower *Foxp3* expression.

These observations underscore a potential shortcoming of unbiased clustering distinguishing Tconvs from Tregs but more importantly, highlight the potential implications of a singular master transcription factor difference despite strong phenotypic overlap between rejection and tolerance phenotypes. Whether the Foxp3^+^ cells within the Tfh and non-Tfh clusters are Tconvs transitioning to becoming iTregs, or Tregs transitioning into ex-Tregs, requires further studies^42^.

The divergent functional outcomes to a semi-allogeneic fetus versus transplanted organs led us to investigate the transcriptomes of 2W:I-A^b^ CD4^+^ T cells in naive pregnancy. We observed a modest expansion of effector and regulatory *CD44^+^* cells, with 4 of 8 clusters retaining antigen-inexperienced markers: *Sell* and *Satb1*. These data highlight the divergent transcriptional responses in CD4^+^ cells to allogeneic conflict in pregnancy and transplantation. Pregnancy prevented the full differentiation of Tfh and non-Tfh effector clusters observed in transplant rejection but allowed the modest expansion a *Cd44^+^* effector cluster, E2, that was remarkably similar within the cluster from naive virgin mice. Importantly, this E2 cluster lacked an *Ifng^+^* signature, but instead expressed regulatory markers such as *Foxp3*, *Areg, and Ctla4,* recapitulating the E1 cluster from transplant tolerance. This suggests a potentially conserved *Foxp3^pos^* mechanism in tolerance across pregnancy and transplantation. We anticipate that the articulation of these atypical effector CD4^+^ T cells expanded in pregnancy and transplant tolerance may pave the way for their characterization in other contexts of immune hypofunction.

Successful parturition in rejection-sensitized dams requires not only the control of antigen-specific naive but also memory T cells. Focusing on 2W:I-A^b^-specific CD4+ T cells, we showed that skin-sensitized mice were dominated by an *Il7r^+^* memory cluster, while a compendium of effector Tconvs and Tregs was observed in SPg. Effector-like non-Tfh and Tfh clusters were induced in sensitized pregnancy with the Tfh-like clusters confirmed by protein expression with Radviz visualization showing IL21^+^ and IL21^-^ polarization of Tconvs in both spleen and lymph nodes. Importantly, SPg E2 and Tfh clusters were transcriptionally enriched for gene sets expressed by transplant Tx E1-3 and F1-3 clusters, as well as for ScRNA gene sets of human graft-infiltrating Tfh-like cells from kidney transplant rejection^31Shi^. The emergence of Tfh clusters was also congruent with the anti-2W-OVA IgG response in Spg and Rej. Additional sub-analysis of pooled follicular T cells from Rej and SPg revealed a Foxp3^+^Tfr cluster present in both, and a Foxp3^neg^ TR1-like cluster observed only in SPg^36, 43^. Indeed, TR1 cells have recently been reported in multiparous mice, suggesting a conserved mechanism in sensitized pregnancy to constrain memory T cell responses^44^.Whether Tfr populations emerge in SPg from Tregs or Tfh, and constrain Tfh responses by inhibiting T effector cells or promoting B cell differentiation into Bregs, requires further investigation ^43, 45, 46, 47, 48^. While the non-Tfh effector clusters in sensitized pregnancy also enriched transcriptionally for transplant E2, it is notable they did not express *Ifng.* Indeed, the absence of *Ifng*-expressing non-Tfh clusters is a notable feature shared with pregnancy in non-sensitized dams and transplant tolerance.

The constraint of mature, effector CD4^+^ T cells is achievable through cell-extrinsic mechanisms exerted by Foxp3^+^ Tregs or other regulatory cells ^49^ while cell-intrinsic mechanisms can be due to anergy or exhaustion, with anergy arising from TCR priming without co-stimulation while exhaustion is induced by persistent antigen exposure ^50^. Transcriptional analysis of these two hypofunctional states have identified heterogenous phenotypes without coalescing to a conserved signature. Early studies by *Sundstedt et al* implicated reduced AP-1 (activator protein 1) and inactive NF-kB complexes in anergic CD4+ T cells ^51^. In contrast, *Trefzer et al* reported that anergic FR4+CD73+ cells co-expressed exhaustion markers such as Lag3, PD1, and CD200^50^. Recent ScRNA-seq studies on oral tolerance reported an expanded endogenous polyclonal antigen-specific CD4^+^ T cells with a FR4^pos^CD73^pos^ Th-lineage-negative (Thlin^-^) anergic subset ^52Hong^. Our studies identified two (E1 and E2) clusters in SPg, and an E2 cluster in Pg that also did not upregulate the lineage-defining effector genes, *Tnf, Ifng*, *Bcl6, Cxcr5* or *Il21*. These clusters instead upregulated *Izumo1r, Nt5e, Ctla4, Nrp1, Ikzf2* and *Tnfrsf18* co-expressed by Tfh and *Foxp3^+^*Treg clusters. However, by trajectory and RNA velocity analyses, these SPg E1-3 and Tfh clusters appeared along different branches with divergent velocity directionality suggesting they are not converging into a common phenotype. Whether these atypical effectors identified in Pg and SPg represent the elusive anergic-exhausted CD4^+^ T cells from previous studies remains unclear. Additionally, their relationship to the Foxp3^+^ E1 cells in transplant tolerance requires further investigation, given that anergic CD4^+^ T cells have been reported to differentiate into regulatory T cells with suppressive function ^12,53,52^.

Observations of Foxp3^+^ Tregs playing critical roles in tolerance across transplantation and pregnancy ^7, 8, 9, 11, 12, 13^ prompted a deeper analysis of Foxp3^+^ cells concatenated across all models revealing substantial Treg heterogeneity. Central Tregs, most represented in our naive groups overlapped with published quiescent mouse and human Tregs. All other Treg clusters were strongly represented in SPg, suggesting a diverse regulatory landscape uniquely induced by pregnancy in a backdrop of memory antigen-specific CD4^+^ T cells generated by skin-graft rejection. Effector-Low Tregs expressed several S100 family proteins as top upregulated DEGs; S100 proteins are responsible for signaling cascades of immune functions including motility, inflammation, and differentiation ^54^. The increasing *Il10* expression in the Proliferating and Effector-Low Tregs suggests that they are maturing into Effector-High Tregs. Consistent with this, trajectory analysis revealed a shared branch spanning Proliferating, Effector-Low and Effector-High Tregs. Interestingly, while the *Bcl6^+^Cxcr5^+^* Follicular, Effector-Low and Effector-High Tregs co-expressed genes mediating regulation, *Tgfb1, Nt5e* and *Areg,* follicular Tregs did not upregulate effector genes (*Il10, Entpd1, Klrg, Itgae, Gzm*b) unique to Effector-High Tregs^55^. These observations underscore discrete effector versus follicular Treg clusters that mirror the phenotypes of Tconvs observed in both transplantation and pregnancy.

Finally, we identified a *Foxp3* High cluster prominently expanded in transplantation with robust expression of *Foxp3, Ikzf2, and Tgfb1* and several splicing genes as top upregulated DEGs. Indeed, throughout our transplant and pregnancy models we consistently identified an “Unclassified” cluster expressing the genes *Srrm2* and *Lars2* involved in RNA splicing of immature into mature RNA, and protein synthesis through tRNA charging and mTORC1 activation, respectively ^26, 28, 56^. We speculate that the *Foxp3* High is a transitioning cluster based on its enrichment for metabolism of lipids and phospholipids^57^. Consistent with this hypothesis, *Itga4*, a migration associated molecule ^58^, and *Klrg1,* which were highly expressed in the Effector High Treg cluster, had positive RNA velocity in the Foxp3 High cluster suggesting increased nascent mRNA for these markers. Deciphering the trajectory and phenotype of effector Tregs will be useful in better articulating the roles in disease pathology.

Importantly, we show conserved signatures of mouse antigen-specific Tregs with published human Tregs. Namely, our proliferating Tregs corresponded with a proliferating Treg gene set in healthy human donors and GVHD patients. Furthermore, our Effector-Low and -High Tregs corresponded with human effector Tregs from uterine tissues in healthy pregnancy. By protein and Radviz visualization, we confirmed Ox40 as a key polarizing marker of activated Tregs in both spleen and lymph nodes across both pregnancy models. Finally, the Treg clusters in successful mouse pregnancy were not enriched for the gene set derived from Early Pregnancy Failure patients with a disrupted Treg transcriptional signature. These observations illustrate the utility of our phenotypic approach in identifying Tregs signatures with sufficient granularity to classify Tregs across species in health and disease.

In its totality, our ScRNA analysis across mouse models of transplantation and pregnancy reveals the extent of heterogeneity in antigen-specific CD4^+^ Tconvs and Tregs. Specifically, our analysis unveiled two bifurcating Tfh and *Ifng*^+^ effector clusters in transplant rejection, and the unexpected co-expression of *Foxp3* by some of these clusters in tolerance. In contrast, naive pregnancy limited T cell effector expansion, while sensitized pregnancy resulted in parturition despite the expansion of Tfh-like clusters and other effector-like clusters. Foxp3^+^ Treg expansion and differentiation was observed across all models, including a TR1 cluster unique to sensitized pregnancy which may be involved in controlling antigen-specific memory responses. In transplantation, overcoming memory to desensitize patients for better transplant outcomes is an active area of research which may be informed by insights from pregnancy in sensitized dams^59^. Regardless of whether these clusters represent temporary effector states along an effector continuum or stable discrete states, we anticipate the utility of these phenotypes lies in their immunosurveillance potential for predicting the outcome of pregnancy and transplant rejection risk. Beyond transplantation and reproductive immunology, we anticipate the T cell biomarkers highlighted in this study will be leveraged in other settings involving adaptive immune responses.

### Limitations of the Study

Pregnancy is a complex, carefully orchestrated biological process involving various biological systems with distinctly described phases ^60^, but we restricted our study to one portion of this process, namely, immediately after parturition. Similarly, transplantation in both tolerance and rejection is marked by distinct phases (early vs late) that can affect T cell phenotypes, but we restricted our study to POD30 when circulating anti-CD154 used to induce tolerance is a fraction of the administered dose. In addition, we did not interrogate graft-infiltrating and fetal/maternal interface lymphocytes but focused on peripheral cells pooled from spleen and lymph nodes, which allowed us to make broad comparisons with T cells from human blood in health and disease. Despite pooling multiple mice for each experimental condition in each model, the recovery of low cell numbers could lead to sampling error in some groups. While low recovery is inherent in profiling an endogenous, antigen-specific population, the ScRNA-seq approach nevertheless allowed us to articulate the transcriptome on a per cell basis across transplantation and pregnancy. Finally, while we highlighted multiple populations, we did not perform functional analyses to verify whether our profiled clusters were functionally regulatory, dysfunctional, or rejecting in a transplant or pregnancy context. Instead, we leveraged canonical markers from the literature to annotate clusters and the models themselves, where the known fate of the transplanted organ or success of the pregnancy, allowed us to generate hypotheses of their function. Future studies will focus on validating the function and interplay of the clusters we articulated as well as their interactions with other cell types.

## ACKNOWLEDGEMENTS

We acknowledge the Cytometry and Antibody Technology Facility (RRID: SCR_017760) at University of Chicago, which receives financial support from the Cancer Center Support Grant (P30CA014599), the Single Cell Immunophenotyping Core and the Animal Resource Center (RRID:SCR_021806). We also thank the NIH Tetramer Core Facility for providing all the tetramers used in this study (contract number 75N93020D00005). This work was supported by grants from the NIH (P01AI097113, R01AI182097) to ASC, MLA and PTS.

## AUTHOR CONTRIBUTIONS

MSA designed all tissue-processing and antibody-staining protocols and conceptualized, constructed, scripted all ScRNA-seq libraries and bioinformatics analysis pipelines. MSA, GH, and ZS prepared mice, processed tissue, and performed flow cytometry experiments. DY performed heart and skin transplants. MLA and PTS provided critical feedback on data analysis. MSA and ASC designed the study, generated the figures, and cowrote the manuscript. All authors edited the manuscript.

## MATERIALS AND METHODS

### Mice

C57BL/6 (B6) and BALB/c (B/c) females, ages 7-8 weeks, were purchased from the Jackson Laboratory or Harlan Laboratories. 2W-OVA B/c males were crossed with B6 females to generate 2W-OVA F1 mice which were used as heart transplant donors. 2W-OVA B/c females were used as skin transplant donors. 2W-OVA B/c males were crossed with virgin B6 females to generate naïve post-partum mice. To generate sensitized post-partum mice, Virgin B6 females transplanted with 2W-OVA skin were bred with 2W-OVA B/c males at post-operative D30-60. Mouse experiments were approved by the Institutional Animal Care and Use Committee at the University of Chicago using standards of the NIH *Guide for the Care and Use of Laboratory Animals* (National Academies Press, 2011).

### Tissue Processing

Spleens and inguinal, axial, and brachial lymph nodes were harvested and passed through a 40 μM strainer (Corning, catalog 431750). Lymphocytes were enriched for CD4^+^ (Miltenyi Biotec, catalog 130-104-454) cells by negative selection prior to antibody staining. Samples were stained for flow cytometry using LiveDead NearIR (Invitrogen, Thermo Fisher Scientific) to exclude dead cells. An antibody cocktail was used to exclude unwanted cells, consisting of CD49b (DX5, catalog 485971-82, Invitrogen, Thermo Fisher Scientific), CD11c (N418, catalog 48-0114-82), F4/80 (BM8, catalog 48-4801-82, Invitrogen, Thermo Fisher Scientific), NK1.1 (PK136, catalog 48-5941-82 eBioscience, Thermo Fisher Scientific), Ter-119 (Ter-119, catalog 48-5921-82, eBioscience, Thermo Fisher Scientific), and CD19 (eBio1D3, catalog 48-0193-82, Invitrogen, Thermo Fisher Scientific). Additional antibodies for phenotyping against CD90.2 (53-2.1, 565257, BD Biosciences), CD4 (GK1.5, 612952, BD Biosciences), Foxp3 (FJK-16s, catalog 53-5773-82, Invitrogen, Thermo Fisher Scientific), Bcl-6 (358509, Biolegend), Icos (117427, Biolegend), CD200/Ox2 (123817, Biolegend), Itga4 (103611, Biolegend), CD27 (124233, Biolegend), IL21 (48-7211-82, Thermo), Cxcr5 (752549, BD), CD134/OX40 (119411, Biolegend), Cxcr4/CD184 (146517, Biolegend), Ccr7/CD197 (120127, Biolegend), CD279/PD-1 (135241, Biolegend), and PD-L1 (569606, BD) were used to stain T cells. The NIH Tetramer Facility provided PE and APC-conjugated 2W (EAWGALANWAVDSA):I-A^b^ tetramers. Tetramer stains were incubated for 40 minutes at room temperature prior to the addition of all other antibodies. The FACSAria Fusion (BD) was used for cell sorting or the Cytek Aurora (Cytek Biosciences) was used to quantify flow cytometry experiments. Cells were fixed and permeabilized if intracellular stains were required.

### Donor/Fetal-specific antibody quantification

Donor/Fetal-specific antibodies in the serum of post-partum mice or post-transplanted mice were measured using 1 × 10^6^ 2W-OVA B6 splenocytes which were incubated for 30 minutes at 4°C with 2 μL of serum from recipient mice. Cells were then washed and incubated with anti-CD19 (1D3, catalog 550992, BD Biosciences) and goat anti-mouse IgG (H+L) (catalog 1031-02, Southern Biotech) for 30 minutes at 4°C. MFI of the CD19^−^ cells that were IgG positive was measured by flow cytometry on the Cytek Aurora.

### ScRNA Library Construction and Processing

Single cell suspensions of lymphoid tissue from each biological group were stained with distinct barcoded antibodies (Cell-Hashtag Oligonucleotides, TotalSeq-B, Biolegend) and sorted on CD90.2+CD4+ double 2W+ tetramer positive (PE and APC) cell surface markers at the University of Chicago Flow Cytometry Core. The 10x Chromium was used in the University of Chicago Single Cell Immunophenotyping Core. Single cell 3’ Kit v3.1 Dual Index Gene Expression with Feature Barcoding, 10x Genomics kit was used to generate cDNA, mRNA, and hashtag oligonucleotide (HTO) libraries per manufacturer’s directions. The 3’ mRNA libraries were sequenced to ∼ 40,000 reads per cell while the HTO libraries to at least 5000 reads per cell using a Novaseq 6000 SP Flow cell. Fastq files from bcl files were generated using Cell Ranger mkfastq and alignment was performed using the STAR aligner against the mm10 transcriptome. Cell Ranger outputs were loaded into Seurat for further QC exclusion of cells > 25% mitochondrial reads. Mitochondrial and ribosomal genes were excluded from downstream analysis.

### Bioinformatics

Standard Seurat processing workflow was used with top n=6000 most variable genes used for PCA generation. Unbiased clusters were generated using the FindClusters command with a resolution of 0.7 for all datasets. Uniform Manifold Approximation and Projection (UMAP) was used for dimensionality reduction taking the top n=20 dimensions from the PCA analysis for all datasets. Differentially expressed genes (DEGs) were found using the FindAllMarkers command using default parameters. Module scores were computed using the AddModuleScore function using default parameters. Pseudotime trajectory analysis was performed using the Monocle3 package while Cell-Cell interaction analysis was performed with CellChat using the filterCommunication function with a minimum of n=5 cells. ClusterProfiler was used for Gene Ontology (GO) overrepresentation analysis. AUCell from the SeuratExtend package was used to calculate gene set enrichment analysis scores (GSEA) against custom gene sets as well as the Reactome database with default parameters. Radviz analysis was performed using the Radviz package according to the default workflow. Briefly, MFI data from each cell of flow cytometry fcs files was exported to R and normalized, markers were used to generate anchors, and the cosine similarity was used to optimize the position of the anchors.

### Statistics

Statistical calculations were made using R or GraphPad Prism Version 10.2.3. *P*-values below 0.05 were considered significant. For pairwise comparisons and the FindAllMarkers function, the Wilcoxon rank sum test was used, and the Holm method to adjust p-values was used for multiple pairwise comparisons. Ordinary one-way ANOVA was used for DSA calculation.

---

**Supplemental Figure 1:**
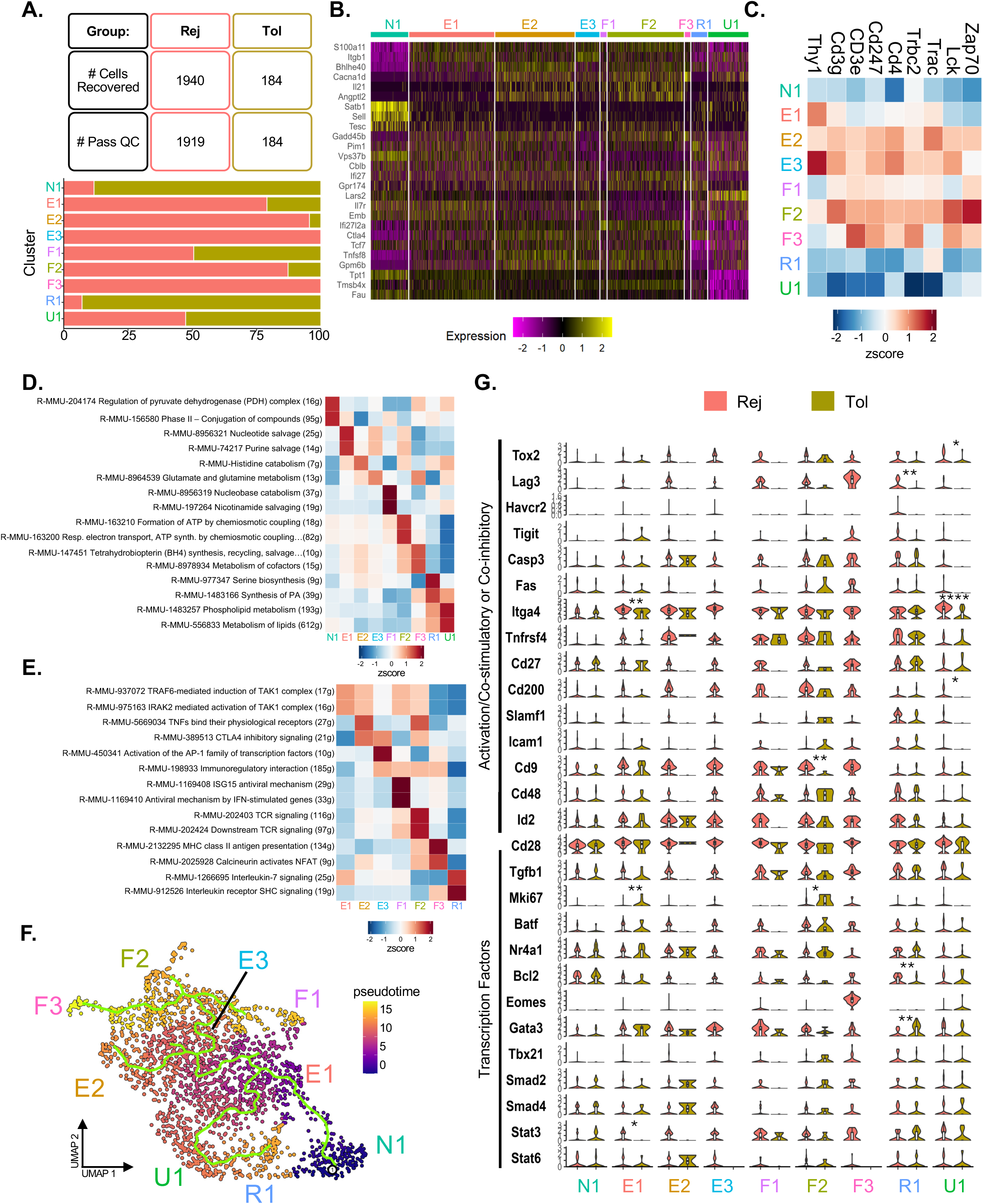
Extended transcriptional analysis of 2W-OVA:I-A^b^ CD4^+^ T Cells from transplant Tol vs Rej. **A.** Total Recovery, number of cells that passed QC, and percentage breakdown of each unbiased cluster by biological group in Rej or Tol 2W-OVA:I-A^b^ CD4^+^ T Cells. Cells pass QC if <25% mitochondrial reads and nFeature_RNA>100. N1, naive 1; E1, effector 1; E2, effector 2; E3, effector 3; F1, follicular 1; F2, follicular 2; F3, follicular 3; R1, regulatory 1; U1, unclassified 1. **B.** Top n=3 downregulated DEGs in each cluster. **C.** Scaled heatmap of T cell receptor components and T cell lineage genes. **D.** Heatmap of clusters by z-score of top enriched pathways from Reactome Database “Metabolism” of Mus musculus. **E.** Heatmap of effector clusters by z-score of top enriched pathways based on Reactome Database “Immune System” of Mus musculus. Color represents strength or expression. **F.** Trajectory and pseudotime analysis overlaid on UMAP. **G.** Violin plot of clusters based on experimental groups, and their expression of select transcription factors and activation/co-stimulatory/co-inhibitory molecules. Wilcoxon signed-rank test with Holm correction; *p < 0.05, **p < 0.01, ***p < 0.001, ****p<0.0001.

**Supplemental Figure 2:**
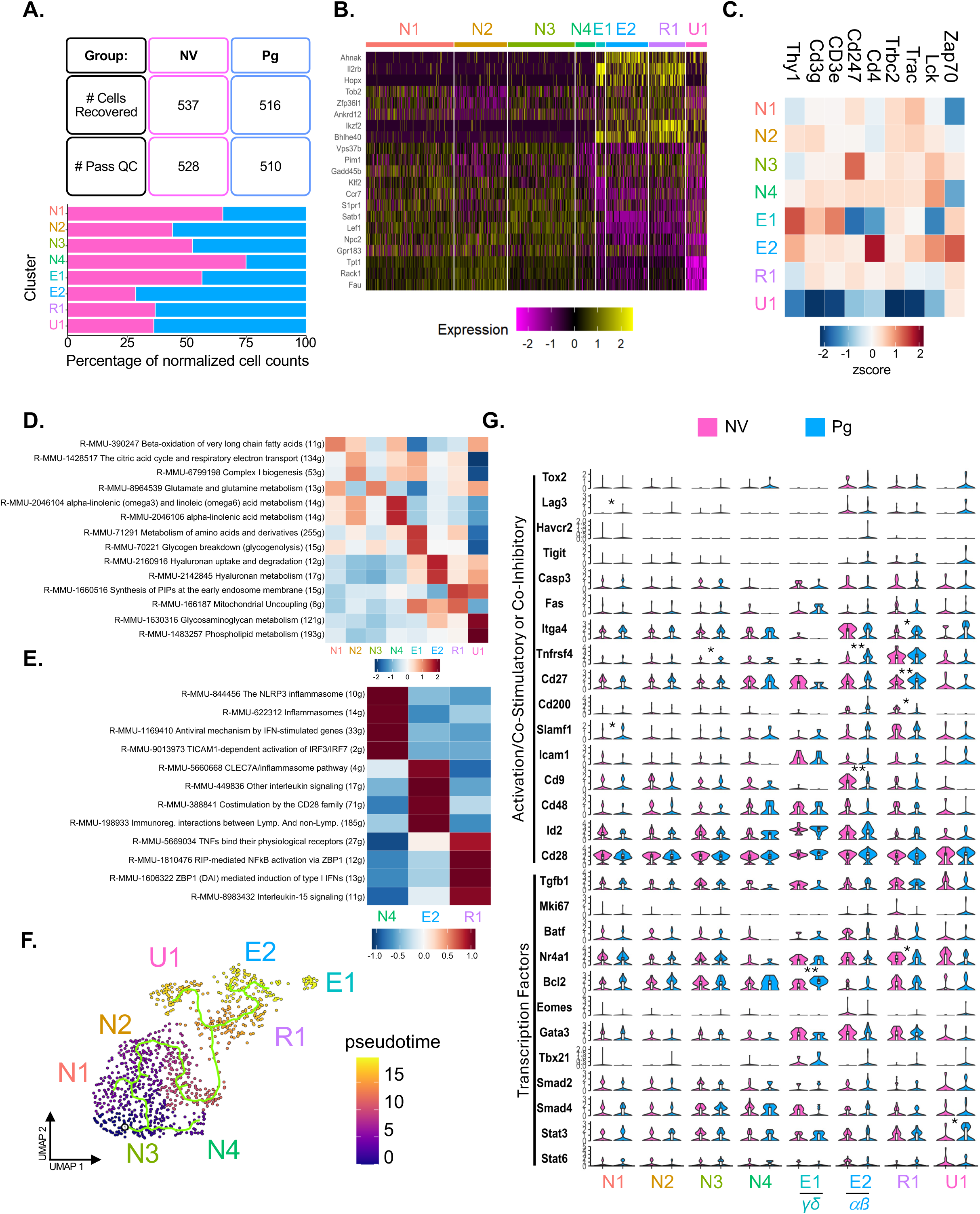
Extended transcriptional analysis of 2W-OVA:I-A^b^ CD4^+^ T Cells from NV vs Pg. **A.** Total Recovery, number of cells that passed QC, and percentage breakdown of each unbiased cluster by biological group in NV or Pg 2W-OVA:I-A^b^ CD4^+^ T Cells. Cells pass QC if <25% mitochondrial reads and nFeature_RNA>100. N1, naive cluster 1; N2, naive cluster 2; N3, naive cluster 3; N4, naive cluster 4; E1, effector cluster 1; E2, effector cluster 2; R1, regulatory cluster 1; U1, unclassified cluster 1. **B.** Top n=3 downregulated DEGs in each cluster. **C.** Scaled heatmap of T cell receptor components and T cell lineage genes. **D.** Heatmap of clusters by z-score of top pathways enriched on Reactome Database “Metabolism” of Mus musculus. **E.** Heatmap of N4, E2 and R1 clusters most modified by pregnancy based on z-scores of top enriched pathways from Reactome Database “Immune System” of Mus musculus. Color represents strength of expression. **F.** Trajectory and pseudotime analysis overlaid on UMAP. **G.** Violin plot of clusters based on experimental groups, and their expression of select transcription factors and activation/co-stimulatory/co-inhibitory molecules. Wilcoxon signed-rank test with Holm correction; *p < 0.05, **p < 0.01, ***p < 0.001, ****p<0.0001.

**Supplemental Figure 3:**
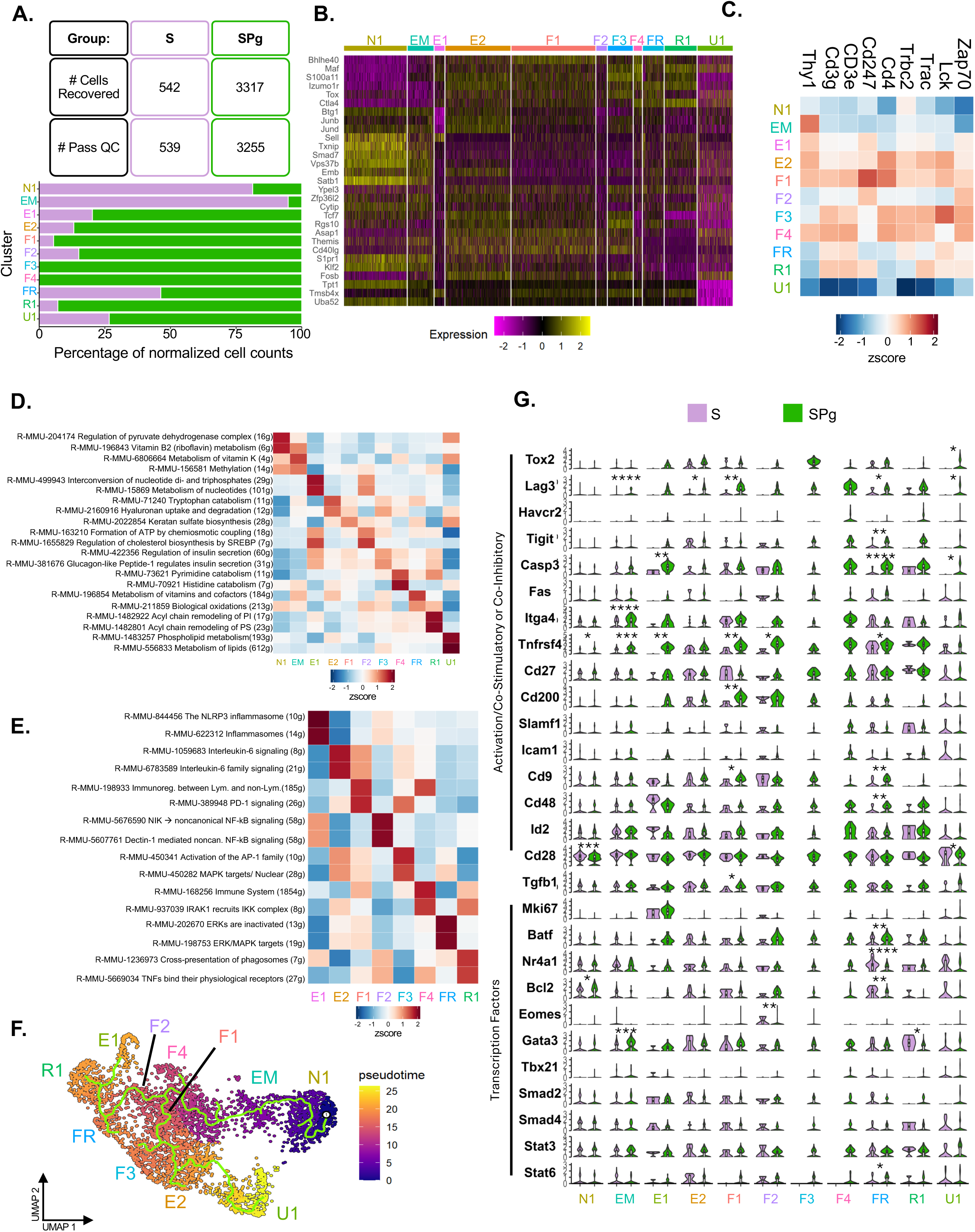
Extended transcriptional analysis of 2W-OVA:I-A^b^ CD4^+^ T Cells from S vs SPg. **A.** Total Recovery, number of cells that passed QC, and cluster breakdown by biological group in S or SPg 2W-OVA:I-A^b^ CD4^+^ T Cells. Cells pass QC if <25% mitochondrial reads and nFeature_RNA>100.). N1, naive 1; EM, effector memory; E1, effector 1; E2, effector 2; F1, follicular 1; F2, follicular 2; F3, follicular 3; F4, follicular 4; FR, follicular regulatory; R1, regulatory 1; U1, unclassified 1. **B.** Top n=3 downregulated DEGs in each cluster. **C.** Scaled heatmap of T cell receptor components and T cell lineage genes. **D.** Heatmap of clusters by z-score of top enriched pathways from Reactome Database “Metabolism” of Mus musculus. **E.** Heatmap of effector clusters by z-score of top enriched pathways from Reactome Database “Immune System” of Mus musculus. Color represents strength of expression. **F.** Trajectory and pseudotime analysis overlaid on UMAP. **G.** Violin plot of clusters based on experimental groups, and their expression of select transcription factors and activation/co-stimulatory/co-inhibitory molecules. Wilcoxon signed-rank test with Holm correction; *p < 0.05, **p < 0.01, ***p < 0.001, ****p<0.0001.

**Supplemental Figure 4:**
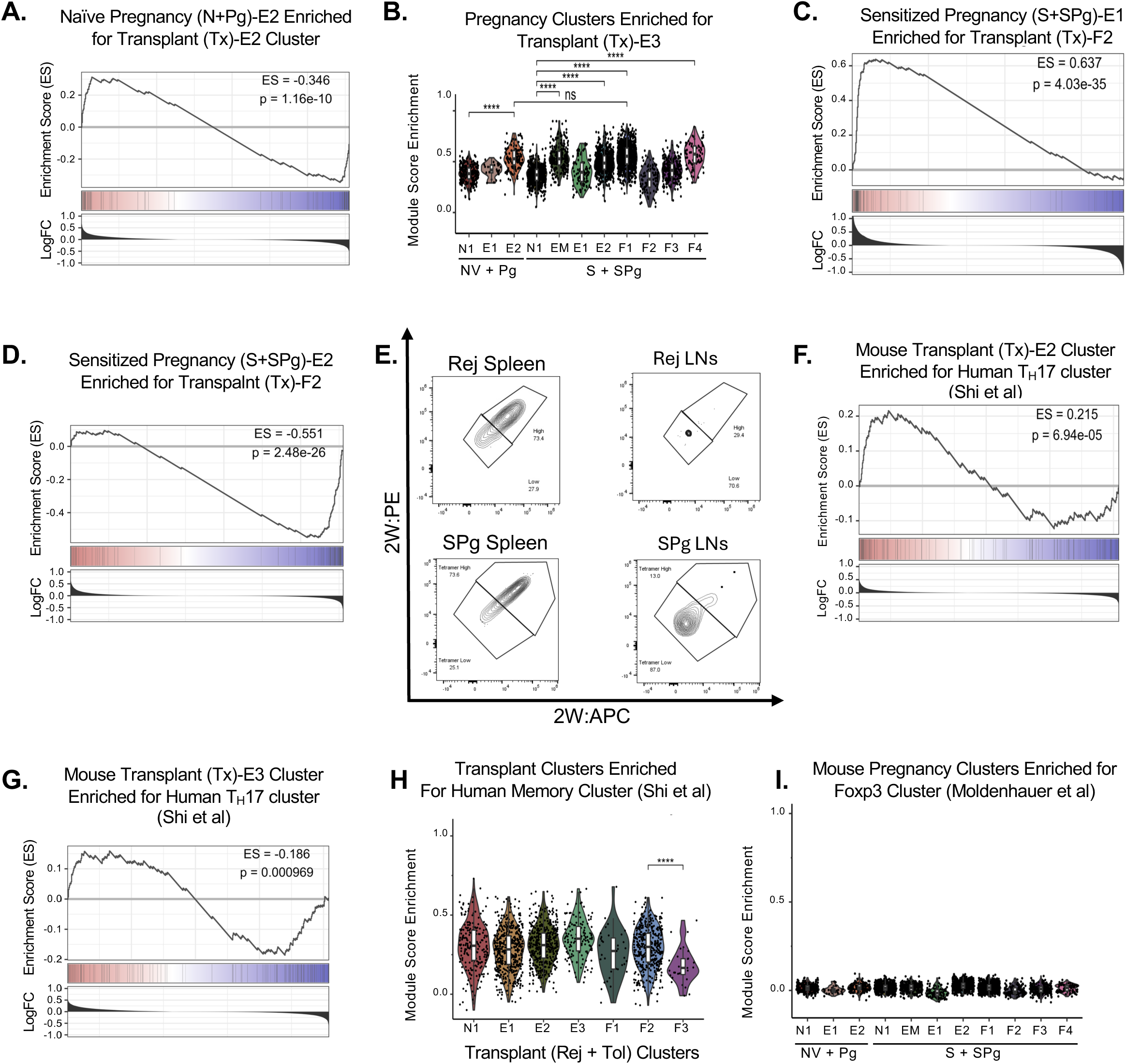
Extended comparative gene set enrichment analysis of *Foxp3*-2W-OVA:I-A^b^ CD4**^+^**transplant and pregnancy clusters for mouse and human datasets. **A.** GSEA of E2 (from NV & Pg) enriched for our Tx-E2 (from Rej & Tol) gene set. **B.** Module score of Naïve pregnancy (from NV & Pg) and Sensitized pregnancy (S & SPg) effector clusters against transplant effector cluster Tx-E3 (from Rej & Tol). Wilcoxon signed-rank test with Holm correction; *p < 0.05, **p < 0.01, ***p < 0.001, ****p<0.0001. **C.** GSEA of E1 (from S & SPg) enriched for our Tx-F2 (from Rej & Tol) gene set. **D.** GSEA of E2 (from S & SPg) enriched for our Tx-F2 (from Rej & Tol) gene set. **E.** Representative flow cytometry plots demonstrate 2W-OVA:I-A^b^ tetramer binding to Tconvs from the spleen or lymph nodes of Rej and SPg. **F.** GSEA of Tx-E2 (from Rej & Tol) enriched for the *Shi et al.* ^31^ “Th17” CD4^+^ gene set. **G.** GSEA of Tx-E3 (from Rej & Tol) enriched for the *Shi et al.* ^31^ “Th17” CD4^+^ gene set. **H.** Module score of transplant clusters (from Rej & Tol) enriched for *Shi et al.* ^31^ “Memory” CD4^+^ gene set. Wilcoxon signed-rank test with Holm correction; *p < 0.05, **p < 0.01, ***p < 0.001, ****p<0.0001. **I.** Module score of Naïve pregnancy (from NV & Pg) and Sensitized pregnancy (from S & SPg) effector clusters against *Moldenhauer et al.* ^32^ “Early Pregnancy Failure TConvs” (EPF) gene set.

**Supplemental Figure 5:**
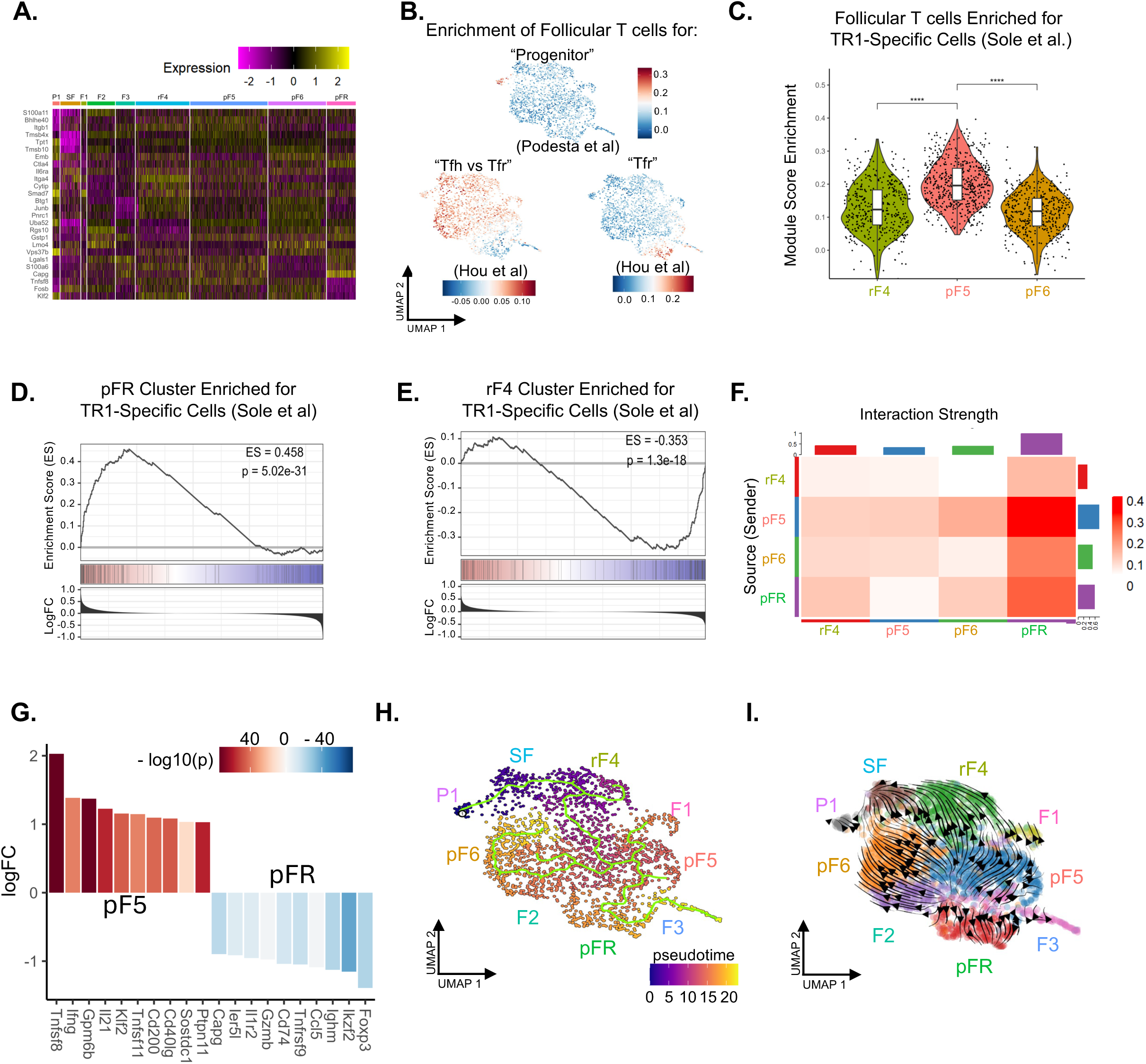
Extended transcriptional analysis, Cell-Cell interaction, Trajectory, and RNA Velocity analysis of Follicular 2W-OVA:I-A^b^ CD4^+^ T Cells in Acute Rejection (Rej) and Sensitized Pregnancy (SPg) **A.** Top n=3 downregulated differentially expressed genes (DEGs) by cluster. Color represents strength of expression. **B.** UMAP and Module score against “Progenitor” (*Podesta et al.*^15^), “Tfh vs Tfr” (*Hou et al.*^35^), and “Tfr” (*Hou et al.* ^35^) gene sets. Color represents strength of expression. **C.** Violin plot of module score by select clusters against TR1-Specific gene set from Solé et al. ^36^. Wilcoxon signed-rank test with Holm correction ; *p < 0.05, **p < 0.01, ***p < 0.001, ****p<0.0001. **D.** GSEA of pFR on *Solé et al.* ^36^ TR1-Specific gene set. **E.** GSEA of rF4 on *Solé et al.* ^36^ TR1-Specific gene set. **F.** CellChat Cell-Cell interactions strength by cluster. Color represents interaction strength. **G.** Waterfall plot showing distinct DEGs from pF5 versus pFR. Color represents strength of expression. **H.** Trajectory and pseudotime analysis overlaid on UMAP. **I.** RNA Velocity by clusters overlaid on UMAP.

**Supplemental Figure 6:**
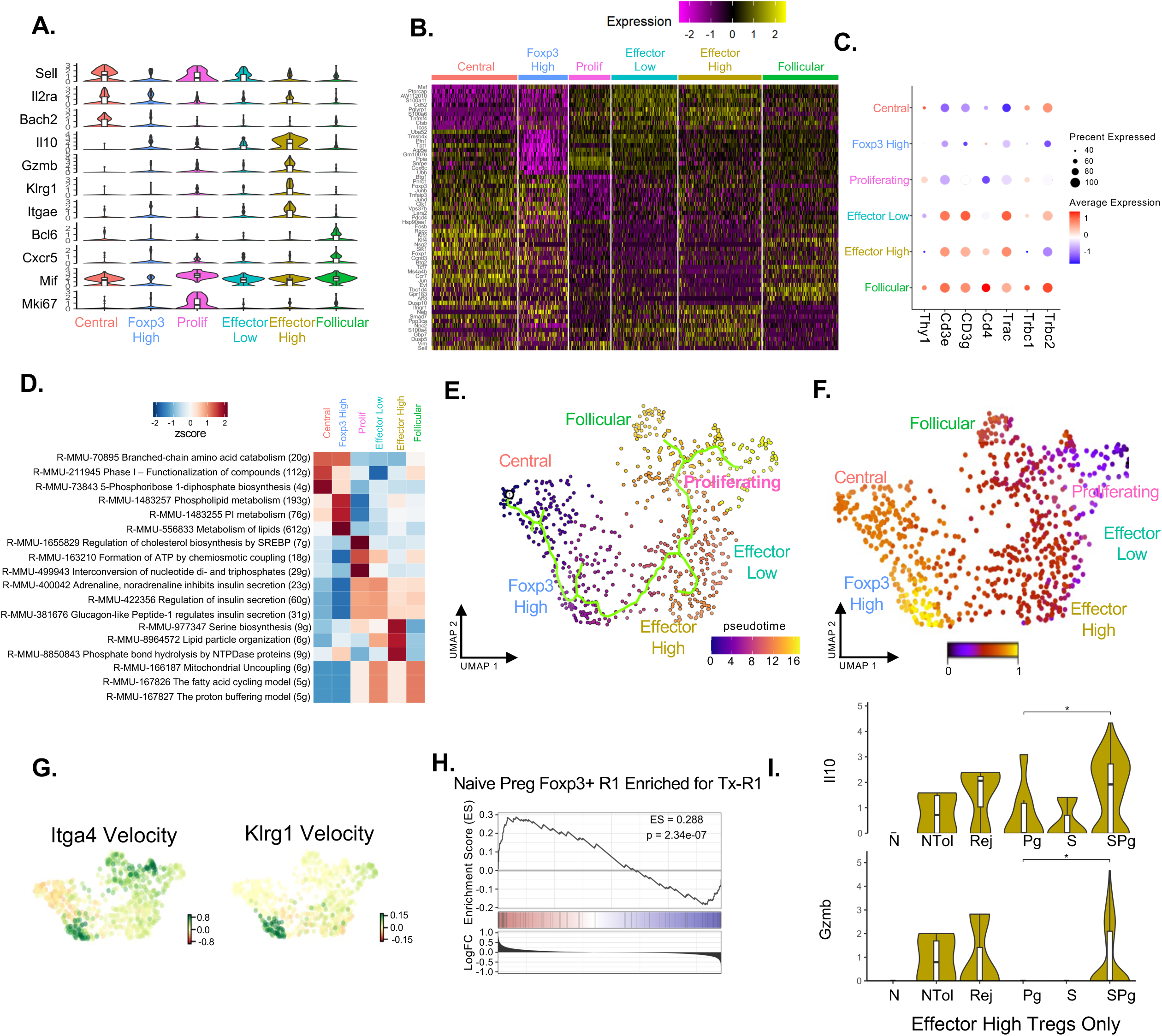
Extended transcriptional analysis, Trajectory, and RNA Velocity analysis of *Foxp3*^+^ 2W-OVA:I-A^b^ CD4^+^ clusters. **A.** Violin plot of markers used for cluster annotations. **B.** Heatmap of top n=10 downregulated DEGs. Color represents strength of expression. **C.** Dot plot of T cell receptor and T cell lineage gene expression by cluster. Size represents percent expression and color represents strength of expression. **D.** Heatmap of clusters by z-score of top enriched pathways from Reactome Database “Metabolism” of Mus musculus. **E.** Trajectory and pseudotime analysis overlaid on UMAP. **F.** Intronic RNA Velocity pseudotime of clusters. **G.** RNA velocity of *Itga4* and *Klrg1*. Color represents upregulation (green) or downregulation (red). **H.** GSEA of R1 (from NV & Pg) versus Tx-R1 (from Rej & Tol). **I.** Violin plots of *Il10* and *Gzmb* of Effector High Tregs split by biological group. Wilcoxon signed-rank test with Holm correction ; *p < 0.05, **p < 0.01, ***p < 0.001, ****p<0.0001.

**Supplemental Figure 7:**
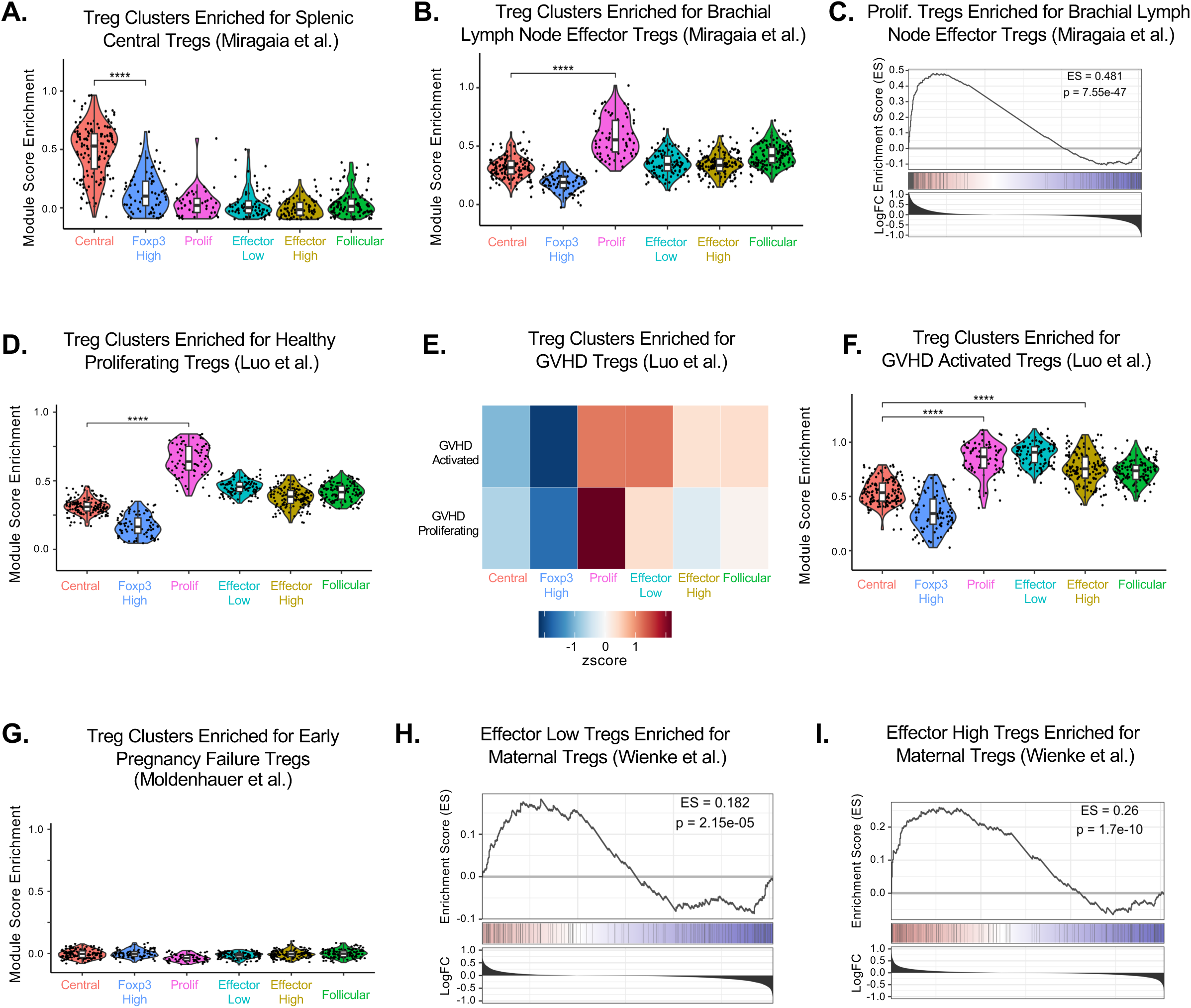
Extended comparative gene set enrichment analysis of transplant and pregnancy *Foxp3*^+^ 2W-OVA:I-A^b^ CD4**^+^** clusters for mouse and human Treg datasets. **A.** Module score of Treg clusters against *“*Splenic Central Tregs” from *Miragaia et al.* ^39^ Wilcoxon signed-rank test with Holm correction ; *p < 0.05, **p < 0.01, ***p < 0.001, ****p<0.0001. **B.** Module score of Treg clusters against *“*Brachial Lymph Node Effector Tregs” from *Miragaia et al.* ^39^ Wilcoxon signed-rank test with Holm correction ; *p < 0.05, **p < 0.01, ***p < 0.001, ****p<0.0001. **C.** GSEA of Proliferating Tregs on *“*Brachial Lymph Node Effector Tregs” from *Miragaia et al.* ^39^ **D.** Module score of Treg clusters against “Healthy Donor Proliferating Tregs” from *Luo et al.*^40^ Wilcoxon signed-rank test with Holm correction ; *p < 0.05, **p < 0.01, ***p < 0.001, ****p<0.0001. **E.** Scaled heatmap by cluster of enrichment for “Graft Vs Host Disease Tregs” from *Luo et al.*^40^ Color represents strength of expression. **F.** Module score of Treg clusters against “GVHD Activated Tregs” from *Luo et al.* ^40^ Wilcoxon signed-rank test with Holm correction ; *p < 0.05, **p < 0.01, ***p < 0.001, ****p<0.0001. **G.** Module score of Treg clusters against “Early Pregnancy Failure TRegs” gene set from *Moldenhauer et al.* ^32^. **H.** GSEA of Effector Low Tregs on *“*Maternal Uterine Tregs” from *Wienke et al.* ^41^ **I.** GSEA of Effector High Tregs on *“*Maternal Uterine Tregs” from *Wienke et al.* ^41^

